# Periostin^+^ stromal cells guide lymphovascular invasion by cancer cells

**DOI:** 10.1101/2022.05.19.492742

**Authors:** Jamie L. Null, Dae Joong Kim, James V. McCann, Patcharin Pramoonjago, Jay W. Fox, Pankaj Kumar, Lincy Edatt, Chad V. Pecot, Andrew C. Dudley

## Abstract

Cancer cell dissemination to the sentinel lymph node associates with poor patient outcomes, particularly in breast cancers. How cancer cells egress the primary tumor upon interfacing with the lymphatic vasculature is complex and driven by dynamic interactions between cancer cells and stromal cells including cancer associated fibroblasts (CAFs). The matricellular protein periostin can distinguish CAF subtypes in breast cancer and is associated with increased desmoplasia and disease recurrence in patients. However, since periostin is secreted, periostin-expressing CAFs are difficult to characterize in situ, limiting our understanding of their specific contribution to cancer progression. Here, we used in vivo genetic labelling and ablation to lineage trace periostin^+^ cells and characterize their function(s) during tumor growth and metastasis. We report that periostin-expressing CAFs are spatially found at periductal and perivascular margins, are enriched at lymphatic vessel peripheries, and are differentially activated by highly-metastatic cancer cells versus low-metastatic counterparts. Surprisingly, genetically depleting periostin^+^ CAFs slightly accelerated primary tumor growth but impaired intratumoral collagen organization and inhibited lymphatic, but not lung, metastases. Periostin ablation in CAFs impaired their ability to deposit aligned collagen matrices and inhibited cancer cell invasion through collagen and across lymphatic endothelial cell monolayers. Thus, highly-metastatic cancer cells mobilize periostin-expressing CAFs in the primary tumor site which promote collagen remodeling and collective cell invasion within lymphatic vessels and ultimately to sentinel lymph nodes.

**Significance Statement:** Metastatic disease causes the majority of cancer-related deaths but is challenging to treat as it is a complex multi-step process driven by heterotypic cell interactions. Cancer-associated fibroblasts (CAFs) are abundant in most solid tumors and display pro-tumorigenic and pro-metastatic functions, but extensive molecular diversity among CAFs has yielded contradictory results in previous attempts to target this population. Therefore, there is a need to identify markers of CAF subpopulations that promote or inhibit metastasis and functionally characterize them to understand their contributions during tumor progression. Our work identifies a population of CAFs, marked by expression of the matricellular protein periostin, that remodel the ECM to promote the escape of cancer cells into lymphatic vessels thereby driving colonization of proximal lymph nodes.

## Introduction

The overwhelming majority of cancer-associated deaths, including those of breast cancer patients, are caused by metastatic burden rather than primary tumor growth. However, the complex process of cancer cell dissemination and colonization of distant tissues is incompletely understood and largely incurable using existing therapies (1). Metastatic disease is particularly intractable because metastasis is not solely driven by cancer cell-intrinsic properties but is instead a consequence of dynamic crosstalk between cancer cells and other cell types in the tumor microenvironment including vascular cells, immune cells, and cancer-associated fibroblasts (CAFs). A critical early step in the metastatic cascade is the intravasation of cancer cells into blood and lymphatic vessels which serve as routes for cancer cells to spread to secondary sites. Lymphovascular invasion of cancer cells into lymphatic vessels is a predominant method of vascular invasion in breast cancer and is significantly associated with the presence of lymph node metastasis, development of distant metastasis, and decreased disease-free interval and overall survival (2-6). Despite the frequency of lymphovascular invasion in breast cancer and its association with poor clinical outcomes, the heterotypic cell interactions that drive lymphovascular invasion are not well characterized. Many studies of lymphovascular invasion focus on paracrine signaling between cancer cells and lymphatic endothelial cells with limited consideration given to the contributions of other auxiliary cells in the tumor microenvironment (TME) such as CAFs.

CAFs are an abundant stromal cell in the breast tumor microenvironment and primarily originate from tissue-resident fibroblasts that become persistently activated by a dysregulated “wound-healing” process recapitulated in solid tumors (7). They are important during multiple stages of tumor development though their specific contributions to disease progression are contradictory and context-dependent (8-14). The paradoxical functions of CAFs during tumor progression can be attributed to their extensive molecular diversity both within and across tumor types as revealed by a number of single-cell RNA sequencing studies (15-21). Though CAFs are mainly characterized by expression of contractile proteins such as alpha smooth muscle actin (SMA) and extracellular matrix (ECM) proteins including collagens and fibronectin, single-cell transcriptomic analysis has identified additional subpopulations of CAFs that display expression profiles associated with proliferation, vascular development and angiogenesis, and immune modulation (15, 22). Although these purported functions of CAF subpopulations position them as attractive therapeutic targets, in vivo depletion studies have yielded unexpected results with SMA^+^ CAF depletion leading to more aggressive, undifferentiated tumors (12, 13). These data suggest that CAF subpopulations play opposing roles in the tumor microenvironment, with some CAFs restraining tumor growth while others are tumor-supportive. Therefore, functional classification of molecularly-defined CAF subpopulations is necessary to overcome the limitations of broadly targeting CAFs so that specific tumor-promoting and/or metastasis-promoting CAFs can be identified. A challenging but critical first step in this process is identifying reliable biomarkers that distinguish pro-tumorigenic and pro-metastatic CAFs from their counterparts.

A recent single-cell RNA sequencing study demonstrated that periostin, a TGFβ-induced matricellular protein, is expressed by CAF populations in human triple negative breast cancers (22). As a nonstructural matrix component, periostin directly interacts with other ECM proteins including lysyl oxidase, collagen I, and fibronectin, serving as a protein scaffold for mediators of collagen cross-linking and ECM stiffening (23). It is typically present at low levels in most tissues but induced at sites of inflammation and tissue injury. Periostin’s role in pathologies including cancer have been noted, including its ability to promote immunosuppressive niche formation, mediate interactions between disseminated cancer cells and the ECM, and support cancer stem cell expansion at metastatic sites (24-26). In human breast cancer samples, periostin is associated with a higher density of activated CAFs and stromal desmoplasia. This stromal desmoplasia is a consistent aspect of invasive progression and distinguishes tumors that are “non-progressors” from those that recur and spread (27). Although these studies characterize periostin as a tumor-supportive factor of metastatic disease, its source in the primary tumor microenvironment and the specific contributions of periostin-expressing cells during tumor progression have not been well-studied in vivo. Previous studies of periostin have relied on immunostaining to characterize its abundance and distribution in primary and secondary sites. However, given that periostin is a secreted factor, immunostaining is unable to link the source of periostin to specific cell types in the tumor microenvironment. Therefore, we have adapted an in vivo lineage tracing strategy previously used to study activated myofibroblasts in the heart (28) to track, characterize, and ablate periostin-expressing cells in the breast TME. We reveal that highly-metastatic cancer cells mobilize periostin-expressing perivascular-like CAFs (PVL-CAFs) in the TME which promote collagen-mediated collective cell invasion of cancer cells across the endothelial barrier of nearby lymphatic vessels and ultimately to the proximal lymph node.

## Results

### Periostin is enriched in CAFs and cycling perivascular-like cells in breast cancers and is associated with advanced disease stage and lymph node metastasis

As periostin has previously been shown to distinguish CAFs in triple negative breast cancer samples (22), we turned to a comprehensive, spatially-resolved single-cell transcriptional atlas of molecularly diverse breast tumors to determine with greater resolution which cellular populations express periostin in human breast cancers (29). We found that periostin is not detected in breast cancer cells of any molecular subtype but is expressed by multiple stromal populations (Figure 1A). Periostin and associated matrix proteins are enriched in CAFs, including mesenchymal stem cell-like inflammatory CAFs (MSC iCAF-like) and myofibroblast-like CAFs (myCAF-like) populations as well as cycling perivascular-like (PVL) cells (Figure 1B). Despite its expression being restricted to the tumor stroma, periostin may have important functional consequences during tumor progression as it is associated with disease recurrence in human breast cancer patients (27). Therefore, we hypothesized that its expression in human breast cancer tissues would correlate with disease stage and lymph node status. Indeed, when we stained a human breast tissue microarray of 200 tumor cores for periostin and measured the positive fluorescent area, we found that periostin abundance positively associated with advanced disease stage (Figure 1C,D) and lymph node metastasis (Figure 1E). These data indicate that periostin is almost exclusively expressed by CAFs or CAF-like cells in the breast TME and its abundance associates with disease progression.

**Figure 1.**
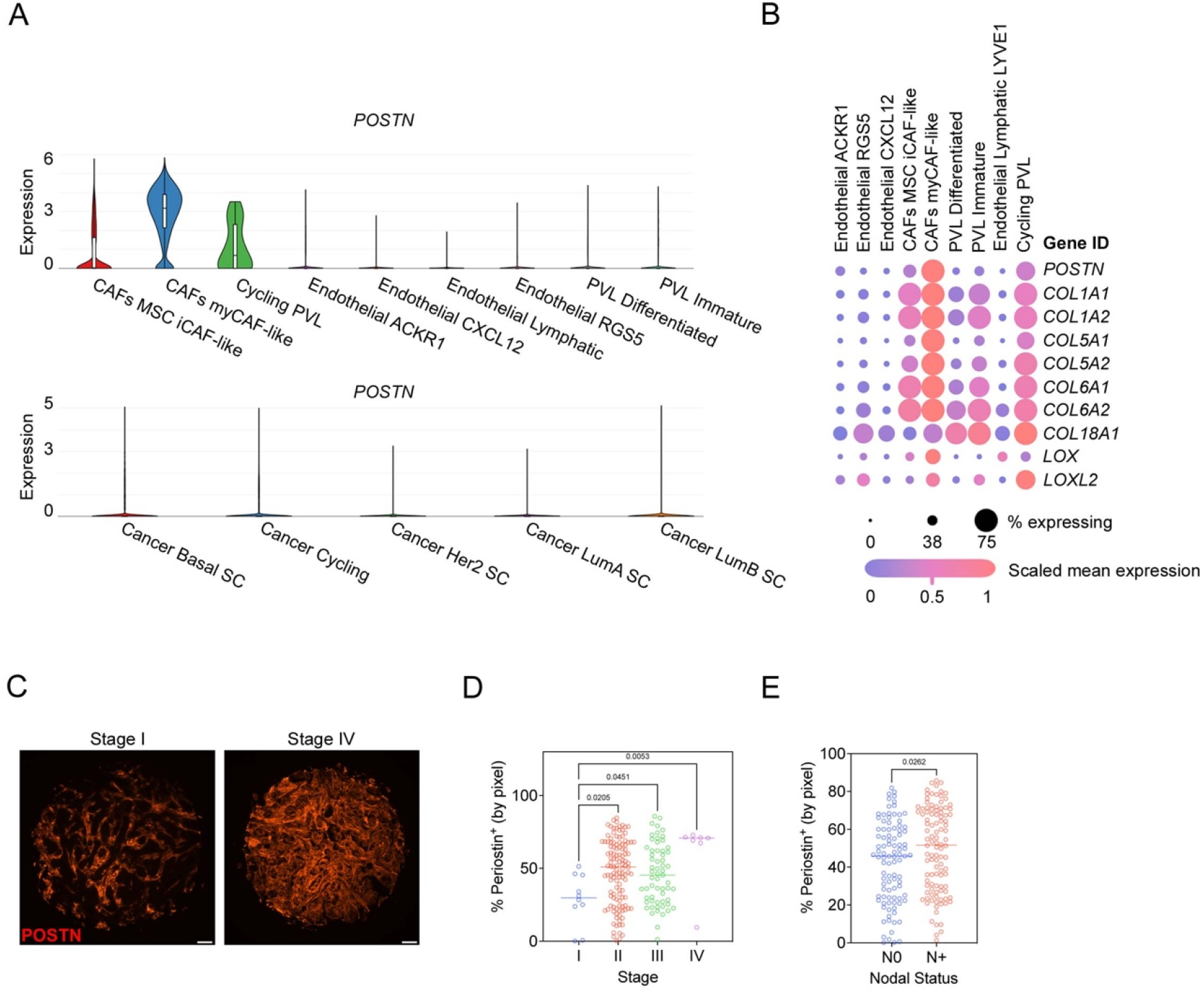
Periostin is enriched in CAFs and cycling perivascular-like cells in breast cancers and is associated with advanced disease stage and lymph node metastasis. (A) Single-cell RNAseq data were downloaded from the Wu et al dataset (29) and analyzed for periostin expression. Violin plots of differential expression of periostin (*POSTN)* in stromal populations (top) and breast cancer cells (bottom) in this patient cohort are shown. MSC = mesenchymal stem cells, PVL = perivascular-like cells, and SC = subclass. (B) Expression of periostin and associated matrix proteins in distinct stromal populations derived from the Wu et al. cohort. (C) Representative immunofluorescence images of human breast cancer tissues stained for periostin (in red). Scale bars: 100 μm. (D) Percentage of tissue area positive for periostin staining (by pixel), grouped by tumor stage (D) and patient’s lymph node status (N0 = no nodal involvement, N+ = nodal involvement) (E). Each data point represents an individual tumor core (N = 200). Statistics shown for ordinary one-way ANOVA, Dunnett’s multiple comparisons test (D) and unpaired Student’s t test (E).

### Periostin-expressing cells surround tumor-naïve mammary ducts and blood vessels and are enriched at the lymphatic vessel periphery

Our human data supports existing murine studies linking periostin to metastasis (24-26). However, previous characterizations of periostin in primary and metastatic environments have primarily relied on immunostaining to characterize its abundance and distribution. Since periostin is a secreted factor, immunostaining is unable to link periostin to a cellular source in the TME. While single-cell RNAseq data indicates that periostin expression is restricted to stromal populations in human breast cancers, these populations have not been studied in situ in murine tumor models. Thus, in vivo approaches to label periostin-expressing cells are required to characterize periostin’s source in the tumor microenvironment and to determine how these periostin-expressing populations function during tumor growth. We generated reporter mice (*Postn*^iZSGreen^ lineage tracing mice) to genetically mark periostin-expressing cells by crossing *Postn*^MCM^ mice with *ZSGreen*^*l*/s/l^ mice. Upon tamoxifen administration, periostin-expressing cells and their progeny are genetically labelled with ZSGreen and can be quantified using fluorescence microscopy which allowed us to spatially track the status of these cells across different tissues. After generating *Postn*^iZSGreen^ mice, we first characterized the spatial distribution of ZSGreen^+^ cells in tumor-naïve mammary glands. We harvested mammary glands from tumor-naïve *Postn*^iZSGreen^ mice and stained for the following cell lineage markers: SMA (fibroblast and pericyte marker), Ck14 (myoepithelial marker), F4/80 (macrophage marker), CD31 (endothelial marker), and Pdpn (lymphatic endothelial marker) (Figure 2A). We found that periostin-expressing cells were relatively sparse in the naïve tissues and surrounded mammary ducts and blood vessels as revealed by the CK14 and CD31 staining, respectively. Intriguingly, we also found that periostin-expressing cells localized to the lymphatic vessel periphery and were more enriched along thin-walled lymphatic vessels compared to large vessel endothelium marked by abundant SMA (Figure 2B), with an average of ∼ one ZSGreen^+^ cell per large blood vessel (± 0.6 cells/vessel) compared to ∼ four ZSGreen^+^ cells per lymphatic vessel (± 0.9 cells/vessel). These data are consistent with the spatially resolved expression of periostin by cycling perivascular-like (PVL) stromal cells in human breast cancers as noted above using comprehensive single cell transcriptomics.

**Figure 2.**
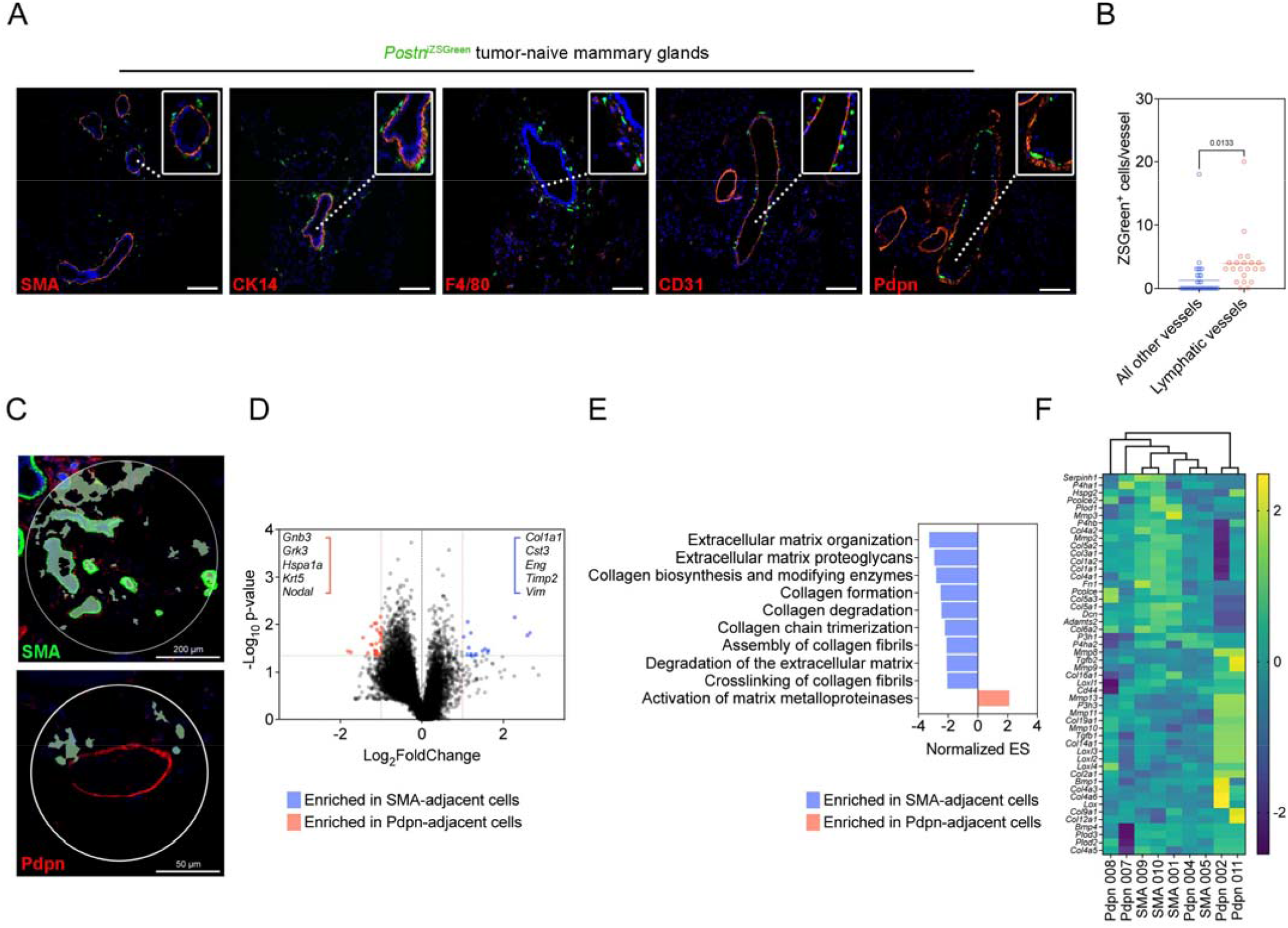
Periostin-expressing cells surround tumor-naïve mammary ducts and blood vessels and are enriched at the lymphatic vessel periphery. (A) Immunofluorescence images of tumor naïve mammary glands from *Postn*^iZSGreen^ lineage tracing mice stained for the following markers (in red): SMA, CK14, F4/80, CD31, and Pdpn. Nuclei counterstained with DAPI. Scale bars: 200 μm, insets are 3x zoom. (B) Quantification of ZSGreen^+^ periostin-expressing cells per vessel (large vessel endothelium marked by abundant SMA versus thin-walled endothelium marked by Pdpn) in serial tissue sections of tumor naïve mammary glands (n = 3 mammary glands per group). Each data point represents an individual vessel. Statistics shown for unpaired Student’s t test. (C) Representative images of regions of interest (ROIs) used to select ZSGreen^+^ periostin-expressing cells for spatial RNA profiling (n = 4-5 ROIs per group, from the mammary glands of 3 mice). (D) Volcano plot of differential gene expression in ZSGreen^+^ cells located near ducts and blood vessels (SMA-adjacent) and ZSGreen^+^ cells located near lymphatic vessels (Pdpn-adjacent). Green line represents a p-value of 0.05 and red line represents the significance threshold. (E) Gene Set Enrichment Analysis (GSEA) showing upregulated pathways in spatially-defined ZSGreen^+^ populations. (F) Cluster analysis of ZSGreen^+^ cells from individual ROIs based on collagen-related genes.

Given the spatially distinct populations of periostin-expressing cells observed along the ducts, blood vessels, and lymphatic vessels in the naïve mammary gland, we used GeoMx Digital Spatial Profiling (DSP) to molecularly characterize the subpopulations in situ and reveal heterogeneity among periostin-expressing cells. Using SMA and Pdpn as morphology markers to select regions of interest within the mammary glands, we performed whole transcriptome analysis on ZSGreen-labelled periostin-expressing cells within these regions (Figure 2C) and compared expression profiles of SMA-adjacent ZSGreen^+^ cells (near ducts and blood vessels) and Pdpn-adjacent ZSGreen^+^ cells (near lymphatic vessels). Spatial transcriptomics identified a number of genes that are differentially expressed between the periostin-expressing subpopulations (Figure 2D), and Gene Set Enrichment Analysis (GSEA) indicated that periostin-expressing cells located near ducts and blood vessels are enriched in genes involved in ECM organization as well as collagen synthesis and remodeling while lymphatic-adjacent periostin-expressing cells are enriched in genes related to the activation of matrix metalloproteinases which degrade the ECM (Figure 2E). However, cluster analysis of the individual subpopulations based on collagen-related genes revealed that there is diversity among lymphatic-adjacent periostin-expressing cells, with multiple lymphatic-adjacent samples enriched in collagens and collagen cross-linking genes suggesting they also play a role in collagen synthesis and organization (Figure 2F). In sum, the spatial transcriptomic analysis demonstrates that these spatially distinct populations of periostin-expressing cells function together to remodel, synthesize, and organize the extracellular matrix within the mammary gland.

### Highly-metastatic mammary tumors differentially activate periostin-expressing PVL-CAFs

Following characterization of periostin-expressing cells in the naïve mammary gland, we next used a mammary tumor model to assess the abundance, morphology, and spatial distribution of periostin-expressing cells in the primary TME and secondary sites of spontaneous metastasis. We used paired, differentially metastatic triple negative mammary cancer cell lines EO771 and EO771.LMB to classify differences in periostin-expressing cell activation in low-versus highly-metastatic tumor contexts. These cell lines have been molecularly and functionally characterized and shown to have differential capacities for spontaneous metastasis, with the poorly metastatic parental EO771 line rarely reaching secondary sites and the highly-metastatic derivative EO771.LMB tropic to the lungs and lymph nodes (30). Importantly, the EO771 and EO771.LMB lines express negligible periostin compared to murine mammary CAFs (Figure 3A), so our lineage tracing strategy successfully captures the predominant host-derived sources of periostin. Following tamoxifen administration to label periostin-expressing cells, *Postn*^*i*ZSGreen^ mice were orthotopically injected with either EO771^mCherry^ or EO771.LMB^mCherry^ cells into the third mammary gland (Figure 3B). According to the previous characterization of the cancer cell lines, the difference in metastatic capacity is observed following primary tumor resection. Therefore, we incorporated this into our study design and harvested secondary tissues 3-4 weeks after tumor resection, allowing more time for disseminated cells to reach and grow out at secondary sites. This experimental setup recapitulates human disease as human breast cancer patients often undergo a lumpectomy but may still progress to metastatic disease as a result of previously disseminated cancer cells emerging from dormancy and expanding in secondary sites. Since the periostin-expressing cells are genetically labelled with ZSGreen and the cancer cells are labelled with mCherry, we could spatially quantify these populations using a thresholding strategy for positive fluorescence in histological sections (Supplemental Figure 1).

**Figure 3.**
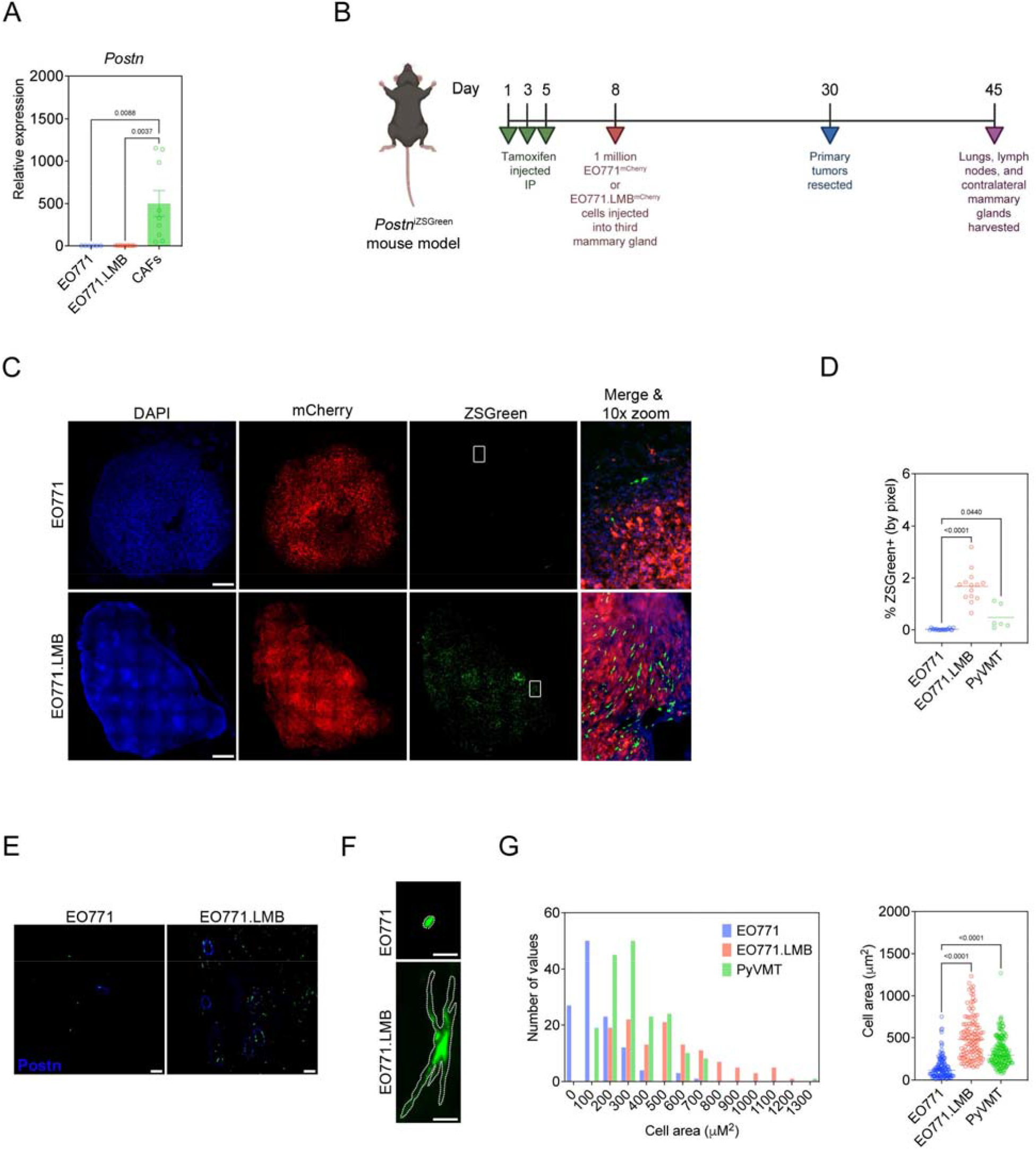
Highly-metastatic mammary tumors differentially activate periostin-expressing PVL-CAFs. (A) qPCR analysis of periostin expression in mammary tumor cell lines EO771 and EO771.LMB and murine mammary CAFs. Data represent the mean ± SEM. Performed in triplicate, and results compared using ordinary one-way ANOVA, Tukey’s multiple comparisons test. (B) Study design including tamoxifen treatment and orthotopic mammary tumor injections for EO771 vs EO771.LMB experiment in *Postn*^iZSGreen^ lineage tracing mice. (C) Tissue tilescans of primary mammary tumors from *Postn*^iZSGreen^ lineage tracing mice. Tumor cells labelled with mCherry and periostin-expressing cells genetically labelled with ZSGreen. Nuclei counterstained with DAPI. Scale bars: 500 μm. (D) Percentage of tissue area positive for ZSGreen in low metastatic (EO771) tumors versus high metastatic (EO771.LMB and PyVMT) tumors (n = 3-7 mice per group). Each data point represents a different histological section. Significance reported in relation to low metastatic (EO771) control, ordinary one-way ANOVA, Dunnett’s multiple comparisons test. (E) Representative immunofluorescence images of mammary tumors stained for periostin (in blue). Scale bars: 100 μm. (F) Representative immunofluorescence images of individual ZSGreen-labelled periostin-expressing cells in low metastatic (EO771) and high metastatic (EO771.LMB) mammary tumors. Scale bars: 10 μm. (G) Area of individual ZSGreen-labelled cells in primary tumors represented as a histogram (left) and scatter plot (right). Each data point in the scatter plot represents an individual cell. Between 120 and 180 cells were measured from multiple tumor sections (n = 3-6 mice per group) and results compared using ordinary one-way ANOVA, Tukey’s multiple comparisons test.

When we examined the low-metastatic EO771^mCherry^ tumors and highly-metastatic EO771.LMB^mCherry^ tumors in the *Postn*^iZSGreen^ mice, we found that periostin-expressing cells were more abundant in highly-metastatic tumors with an ∼ 84-fold increase in the percentage of tissue area positive for ZSGreen in EO771.LMB tumors compared to EO771 tumors (1.68% ± 0.2% versus 0.02% ± 0.01%) (Figure 3C,D). We also examined a second highly-metastatic variant of PyVMT and found a 24-fold increase in the percentage of tissue area positive for ZSGreen in these tumors compared to low-metastatic EO771 tumors (0.48% ± 0.2%) (Figure 3D). This difference in periostin-expressing cell abundance in low-versus highly-metastatic mammary tumors correlated as expected with differences in secreted periostin in situ (Figure 3E). In addition to the difference in abundance of periostin-expressing cells in the low-versus highly-metastatic primary tumors, we observed a difference in the morphology of the ZSGreen^+^ cells between the two groups. Periostin-expressing cells in highly-metastatic EO771.LMB and PyVMT tumors exhibited the typical stellate-shaped morphology of activated myofibroblasts and were 2-3-times larger on average compared to those found in low-metastatic EO771 tumors (509.7 ± 23.6 µm^2^ and 334.2 ± 12.6 µm^2^ versus 146.3 ± 12.0 µm^2^, respectively) (Figure 3F,G). These data suggest that highly-metastatic cancer cells differentially activate periostin-expressing CAFs in the primary tumor compared to their low-metastatic counterparts.

### Periostin-expressing PVL-CAFs are more abundant in the metastatic niches of mice bearing highly-metastatic mammary tumors

Next, we examined the distribution of periostin-expressing CAFs in common metastatic sites of breast tumors, beginning with the draining axillary lymph nodes of the tumor-bearing mice as regional lymph nodes are often the first site of metastatic spread in breast cancers. Periostin-expressing cells were ∼ 14-times more abundant in the axillary lymph nodes of mice bearing highly-metastatic EO771.LMB tumors, with 0.8% (± 0.1%) of the tissue area positive for ZSGreen compared to 0.06% (± 0.03%) in the lymph nodes of mice bearing low-metastatic EO771 tumors (Figure 4A,C). We then assessed the abundance of periostin-expressing cells in the lungs of mice as this is another common site of breast cancer metastasis. Similar to the lymph nodes, we observed a significant increase in the percentage of lungs positive for ZSGreen in mice bearing highly-metastatic mammary tumors compared to those bearing low-metastatic tumors (0.2% ± 0.03% versus 0.08% ± 0.04%) (Figure 4B,D). Interestingly, we observed ZSGreen^+^ cells in a number of histological sections of lymph nodes (Figure 4C) and lungs (Figure 4D) without evidence of mCherry^+^ cancer cells in these same tissues, suggesting that periostin-expressing cells are present in the premetastatic niche. In support of this possibility, we also observed differential activation of periostin-expressing cells in the contralateral mammary glands of tumor-bearing mice, another common metastatic site for breast cancers (31, 32). Though we did not detect mCherry^+^ cancer cells at these sites, there was an ∼ 7-fold increase in the percentage of tissue area positive for ZSGreen in mice bearing highly-metastatic tumors compared to those bearing low-metastatic tumors (1.6% ± 0.2% versus 0.2% ± 0.1%) (Supplemental Figure 2).

**Figure 4.**
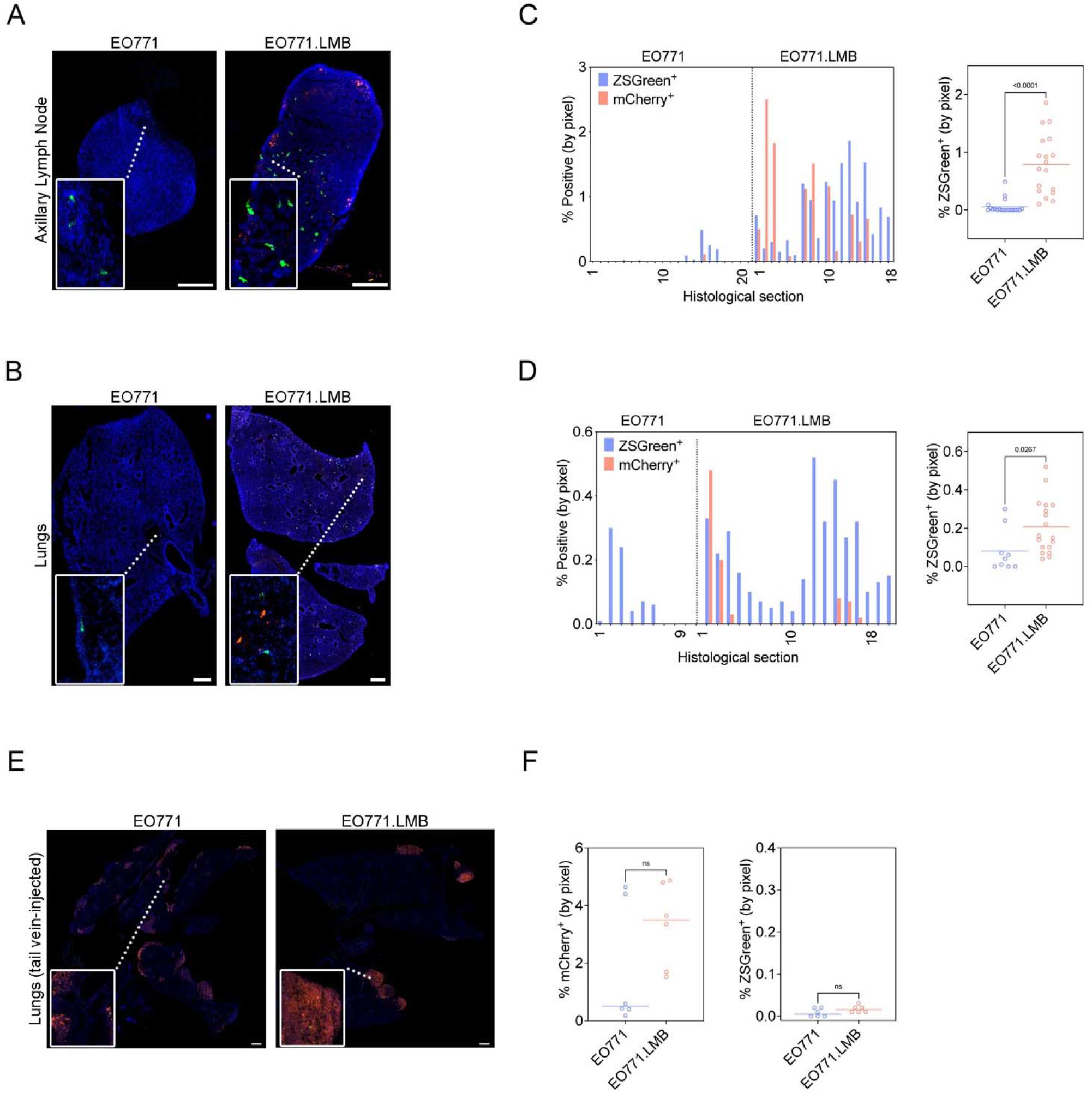
Periostin-expressing PVL-CAFs are more abundant in the metastatic niches of mice bearing highly-metastatic mammary tumors. (A,B) Tissue tilescans of axillary lymph nodes (A) and lungs (B) from *Postn*^iZSGreen^ lineage tracing mice bearing either low metastatic (EO771) or highly metastatic (EO771.LMB) mammary tumors. Tumor cells labelled with mCherry and periostin-expressing cells genetically labelled with ZSGreen. Nuclei counterstained with DAPI. Scale bars: 500 μm, insets are 3x zoom (A) and 6x zoom (B). (C,D) Percentage of tissue area positive for ZSGreen in serial sections of axillary lymph nodes (C) and lungs (D) from mice bearing EO771 or EO771.LMB mammary tumors represented as a histogram (left) and scatter plot (right). Each bar of the histogram represents a different histological section with matched mCherry and ZSGreen measurements. The scatter plot shows individual ZSGreen measurements with each point representing a different histological section (n = 6-7 lymph nodes and 3-6 lungs per group). Statistics shown for unpaired Student’s t test. (E) Tissue tilescans of lungs from *Postn*^iZSGreen^ lineage tracing mice injected via tail vein with EO771 or EO771.LMB mammary tumor cells in a model of experimental metastasis. Tumor cells labelled with mCherry and periostin-expressing cells genetically labelled with ZSGreen. Nuclei counterstained with DAPI. Scale bars: 500 μm, insets are 6x zoom. (F) Percentage of tissue area positive for mCherry (left) and ZSGreen (right) in serial sections of lungs from tail vein-injected mice. Each data point represents a different histological section (n = 3 mice per group). Statistics shown for unpaired Student’s t test.

Since the spontaneous metastases we observed in the lungs resembled micrometastases, we hypothesized that a tail vein model of experimental metastasis would yield greater activation of periostin-expressing cells in the lungs, especially in the EO771.LMB-injected mice, as this would allow a greater number of tumor cells to colonize the lungs and activate periostin expression in tissue-resident cells via growth factor signaling. Surprisingly, though we found macrometastases of both the low-metastatic and highly-metastatic mammary cancer cells in the lungs following tail vein injection (Figure 4E), there was a lack of periostin activation in these tissues (Figure 4F), suggesting that the primary tumor must be established prior to activation of periostin at secondary sites.

### Collagen fibers are longer and more aligned in highly-metastatic (periostin^high^) breast tumors

Because periostin has been shown to promote ECM remodeling by binding to other matrix proteins including collagen, tenascin-C, and fibronectin and enabling collagen cross-linking through a mechanism dependent on BMP-1-mediated activation of lysyl oxidase (LOX) (23, 33), we next used second harmonic generation (SHG) imaging to visualize collagen fibers in low-versus highly-metastatic mammary tumors to determine whether differences in periostin^+^ cell abundance associated with intratumoral collagen structure. Coincident with an increase in periostin-expressing CAFs in highly-metastatic breast tumors, we observed global changes to the collagen matrix architecture in these tumors (Figure 5A). Collagen fibers were more continuous with fewer endpoints in the highly-metastatic tumors (Figure 5B), and these fibers were straighter (Figure 5C) and more aligned when compared to low-metastatic counterparts (Figure 5D). Additionally, there was a 3.7-fold increase in collagen fiber length in highly-metastatic mammary tumors compared to low-metastatic mammary tumors (342.6 ± 14.9 µm versus 93.4 ± 5.4 µm) (Figure 5E). Taken together, these differences indicate that the collagen matrix is more organized in highly-metastatic (periostin^high^) mammary tumors compared to the randomly distributed and shorter collagen fibers found in low-metastatic (periostin^low^) mammary tumors.

**Figure 5.**
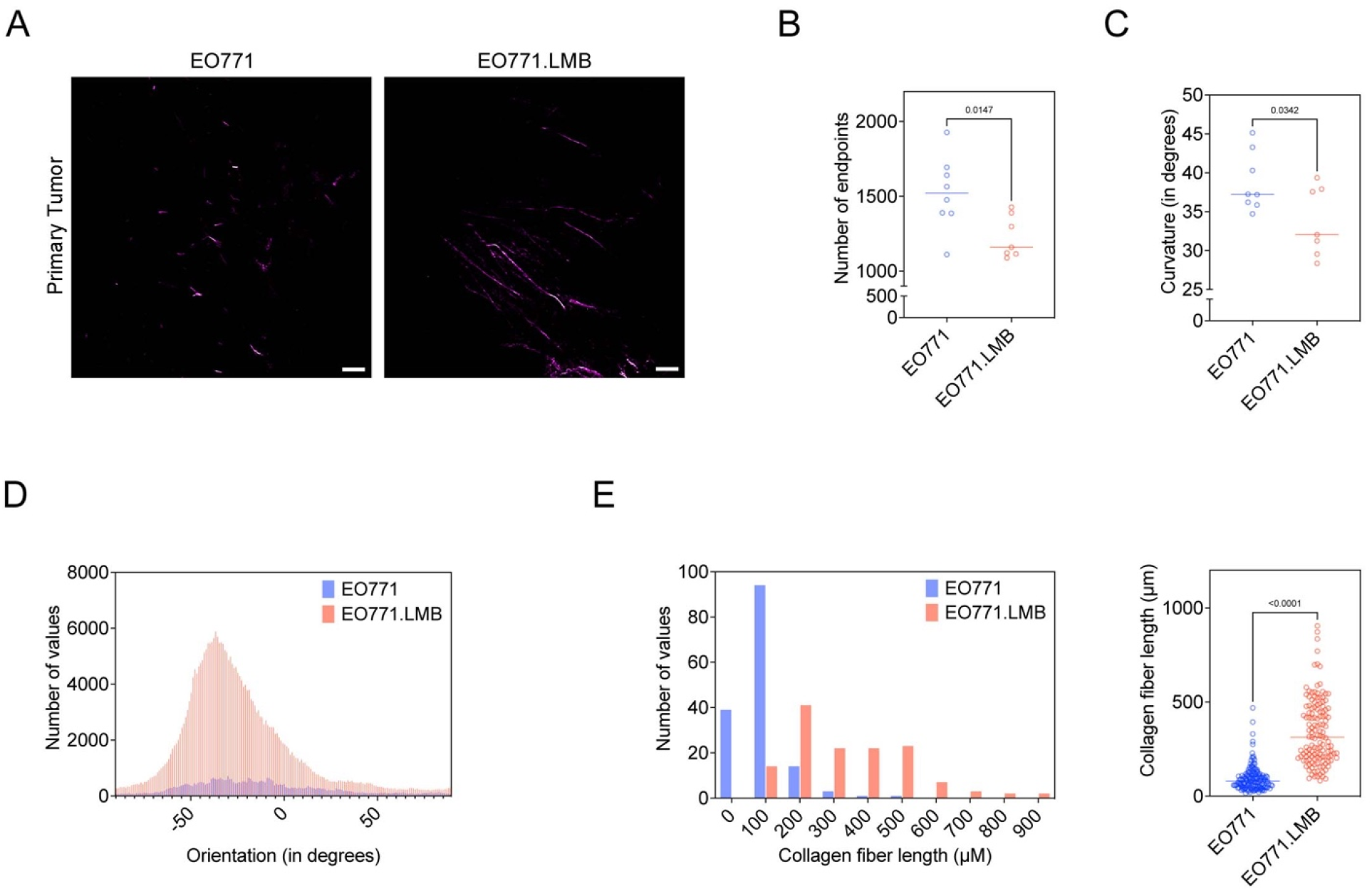
Collagen fibers are longer and more aligned in highly-metastatic (periostin^high^) breast tumors. (A) Second harmonic generation (SHG) images of collagen fibers (pseudo-colored in purple) in EO771 or EO771.LMB mammary tumors. Scale bars: 150 μm. (B) Number of collagen fiber endpoints in histological sections of primary tumors (n = 3-4 mice per group). Each data point represents a different histological section. Statistics shown for unpaired Student’s t test. (C) Curvature of collagen fibers, measured as mean change in angle along fibers, in histological sections of primary tumors (n = 3-4 mice per group). Each data point represents a different histological section. Statistics shown for unpaired Student’s t test. (D) Representative histogram of collagen fiber orientation in primary tumors. A peaked histogram represents aligned fibers whereas a flat histogram represents random organization. (E) Quantification of collagen fiber length in primary tumors, represented as a histogram (left) and scatter plot (right). Each point represents an individual fiber. Between 130 and 160 fibers quantified from multiple histological sections (n = 3 mice per group) and results compared using unpaired Student’s t test.

### Periostin knockdown in primary human breast CAFs alters collagen matrix architecture and inhibits collective cell invasion

Given that an aligned collagen matrix is a signature of more advanced breast tumors and can promote local invasion of cancer cells (34-36), we hypothesized that the difference in collagen matrix structure we observed in vivo was related to differences in the pool of intratumoral periostin and could have consequences for cancer cell migration. Thus, we knocked down periostin in primary human breast CAFs in vitro (Supplemental Figure 3A-C) to determine whether periostin expression itself confers functional properties to CAFs including their ability to deposit an organized collagen matrix. RNAseq of primary human breast CAFs showed that periostin knockdown secondarily reduced expression of matrisomal proteins, ECM regulators, and collagens among other matrix-related genes found in the NABA Gene Set Enrichment Analysis (GSEA) (Figure 6A, Supplemental Figure 3D). Secreted collagen I protein levels were reduced ∼3-fold following periostin knockdown (Figure 6B), and the collagen fibers deposited by periostin-knockdown CAFs were significantly shorter than collagen fibers deposited by CAFs treated with non-targeting control siRNA (201.6 ± 7.6 µm versus 120.0 ± 4.8 µm) (Figure 6C,D). Coupled with our in vivo observations that periostin-expressing cell area and matrix organization were associated with increased intratumoral periostin, these in vitro matrix alterations prompted us to test whether knocking down periostin would affect cell area in vitro, as the stellate-shaped morphology that is characteristic of CAFs is attributed to their ability to engage the ECM and form focal adhesions. Primary human breast CAF cell area was significantly reduced following periostin knockdown (2199 ± 100.2 µm^2^ versus 1643 ± 72.1 µm^2^), and this effect was rescued by addition of recombinant human periostin (2534 ± 131.4 µm^2^) (Figure 6E, Supplemental Figure 3E). The ability of CAFs to spread and form focal adhesions is critical for their motility, so we used migration assays to assess whether periostin knockdown would also impede their ability to migrate. Indeed, periostin knockdown in human breast CAFs reduced migration ∼ 3-fold at 12 hours and 2-fold at 24 hours, and addition of recombinant human periostin restored the migratory capacity of periostin-knockdown CAFs at both time points (Supplemental Figure 3F). Given this observed deficit in CAF migration following periostin knockdown, we hypothesized that ablating periostin in CAFs would also inhibit their ability to promote collective cell invasion. Therefore, we performed a 3D co-culture spheroid assay in which human breast CAFs were treated with either periostin-targeting siRNA (si*-POSTN)* or non-targeting control siRNA (si-Control) and co-cultured in spheroids with MDA-MB-231^mCherry^ human breast cancer cells. CAF/cancer cell spheroids were then embedded in 3D matrices and percent change in invasive area was measured over time. We observed a selective impairment of collective cell invasion through a collagen matrix, as spheroids with periostin knockdown CAFs were significantly less invasive at 24 hours compared with control spheroids when embedded in type I collagen, but there was no difference in invasion when embedded in Matrigel (Figure 6F,G). Taken together, these data suggest that the ability of periostin-expressing CAFs to drive collective cell invasion is selectively dependent on collagen remodeling as there was no invasion deficit following periostin knockdown when spheroids were embedded in Matrigel which primarily consists of laminins and other basement membrane proteins.

**Figure 6.**
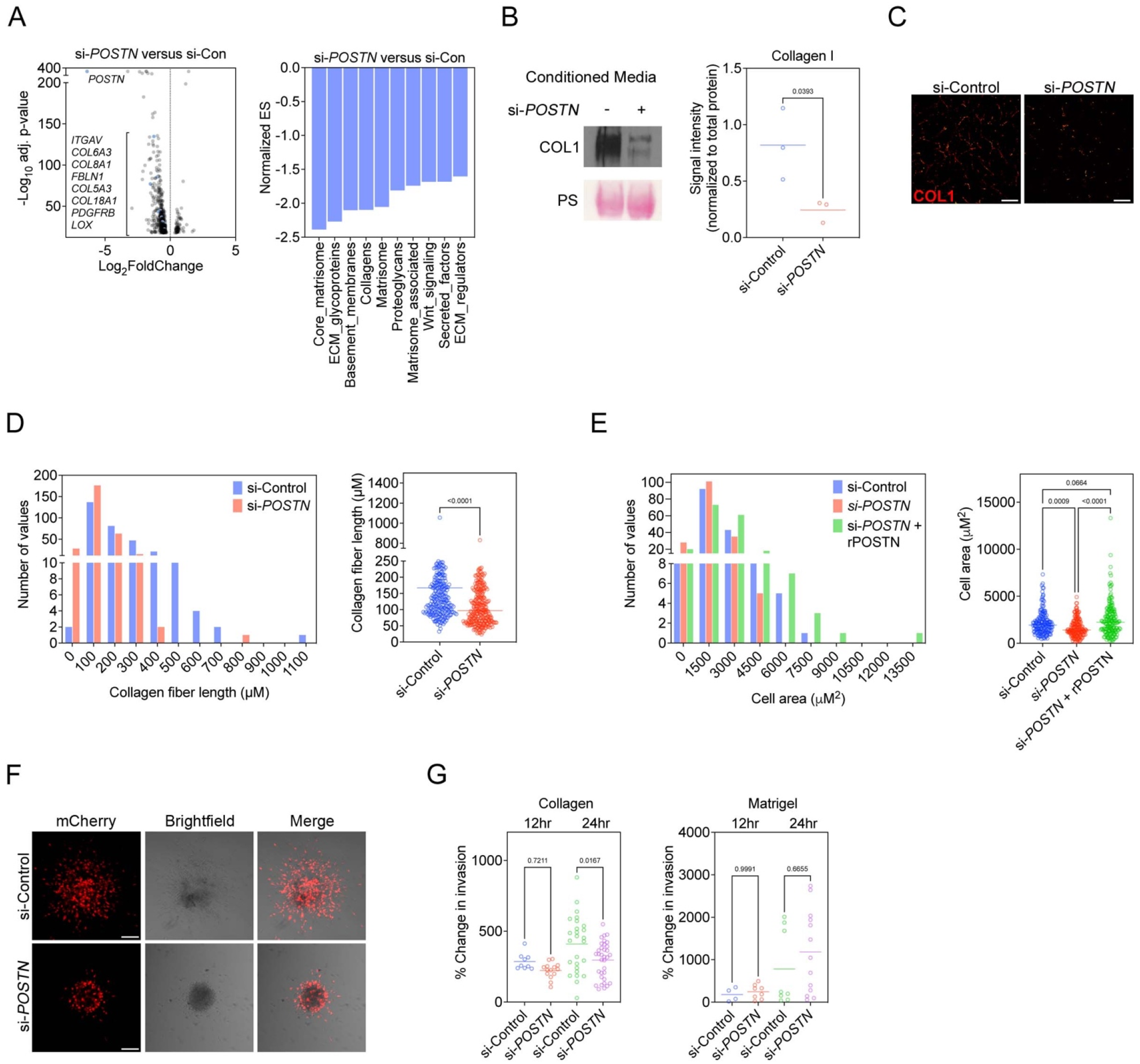
Periostin knockdown in primary human breast CAFs alters collagen matrix architecture and inhibits collective cell invasion. (A) Volcano plot of genes with significantly altered expression detected by bulk RNAseq following periostin knockdown in primary human breast CAFs (left) and NABA Gene Set Enrichment Analysis (GSEA) showing downregulated pathways in periostin knockdown cells (right). Periostin knockdown in human breast CAFs done in quadruplicate. (B) Western blot of secreted collagen I following periostin knockdown. Ponceau stain (PS) shown as loading control. Signal intensity quantified at right and compared using unpaired Student’s t test. Performed in triplicate. (C) Immunostaining of deposited collagen I by si-Control- vs. si*-POSTN-*treated human breast CAFs. Scale bars: 100 μm. (D) Lengths of collagen fibers deposited by control versus periostin knockdown human breast CAFs represented as a histogram (left) and scatter plot (right). Each data point represents an individual continuous collagen fiber (n = 285-305 fibers per group). Experiment performed in triplicate and results compared using unpaired Student’s t test. (E) Cell area measurements of phalloidin-stained primary human breast CAFs treated with si-Control or si*-POSTN* ± recombinant human periostin (rPOSTN) represented as a histogram (left) and scatter plot (right). Experiment performed in triplicate with each data point representing an individual cell (n = 160-190 cells per group). Statistics shown for ordinary one-way ANOVA, Tukey’s multiple comparisons test. (F) Confocal images at the 24 hr timepoint of tumor cell/CAF co-culture spheroids consisting of MDA-MB-231^mCherry^ human breast cancer cells and unlabeled primary human breast CAFs embedded in type I collagen. Scale bars: 100 μm. (G) Percent change in invasive area of tumor cell/CAF co-culture spheroids embedded in type I collagen (left) or Matrigel (right). Each data point represents an individual spheroid, done in biological triplicate. Statistics shown for ordinary one-way ANOVA, Tukey’s multiple comparisons test.

### Periostin-expressing CAFs promote lymphatic metastasis by remodeling the extracellular matrix and directing lymphovascular invasion along organized collagen fibers

Taken together, our in vitro data implicated periostin-expressing CAFs in collagen-mediated invasion. To further explore the specific role of this population during tumor progression and metastasis, we used a DTA depletion strategy to ablate periostin-expressing CAFs in vivo. We generated *Postn*^DTA^ mice by crossing *Postn*^iZSGreen^ lineage tracing mice with *Rosa*-DTA mice. In this model, tamoxifen administration drives expression of diphtheria toxin in periostin-expressing cells resulting in their selective ablation. To confirm ablation of periostin-expressing cells in our *Postn*^DTA^ mouse model, we quantified ZSGreen^+^ cells in the primary mammary tumors of control (*Postn*^iZSGreen^) mice compared to periostin^+^ cell-depleted (*Postn*^DTA^) mice. There was a 75% reduction in the percentage of tissue area positive for ZSGreen in the mammary tumors of the *Postn*^DTA^ mice (1.68% ± 0.2% versus 0.41% ± 0.1%), indicating successful ablation of the majority of periostin-expressing cells (Supplemental Figure 4). We then used these *Postn*^DTA^ mice in our mammary tumor model to measure changes in matrix architecture and metastasis following ablation of periostin-expressing cells (Figure 7A). *Postn*^DTA^ mice were treated with either tamoxifen to induce periostin^+^ cell depletion or vehicle-only control, then injected with highly-metastatic EO771.LMB^mCherry^ mammary cancer cells. Primary tumors were resected and a booster dose of tamoxifen (or vehicle-only control) was administered to ablate any periostin-expressing cells that may have been activated as a result of surgery as periostin is induced by tissue injury and inflammation. Finally, secondary sites were harvested and evaluated for metastatic burden.

**Figure 7.**
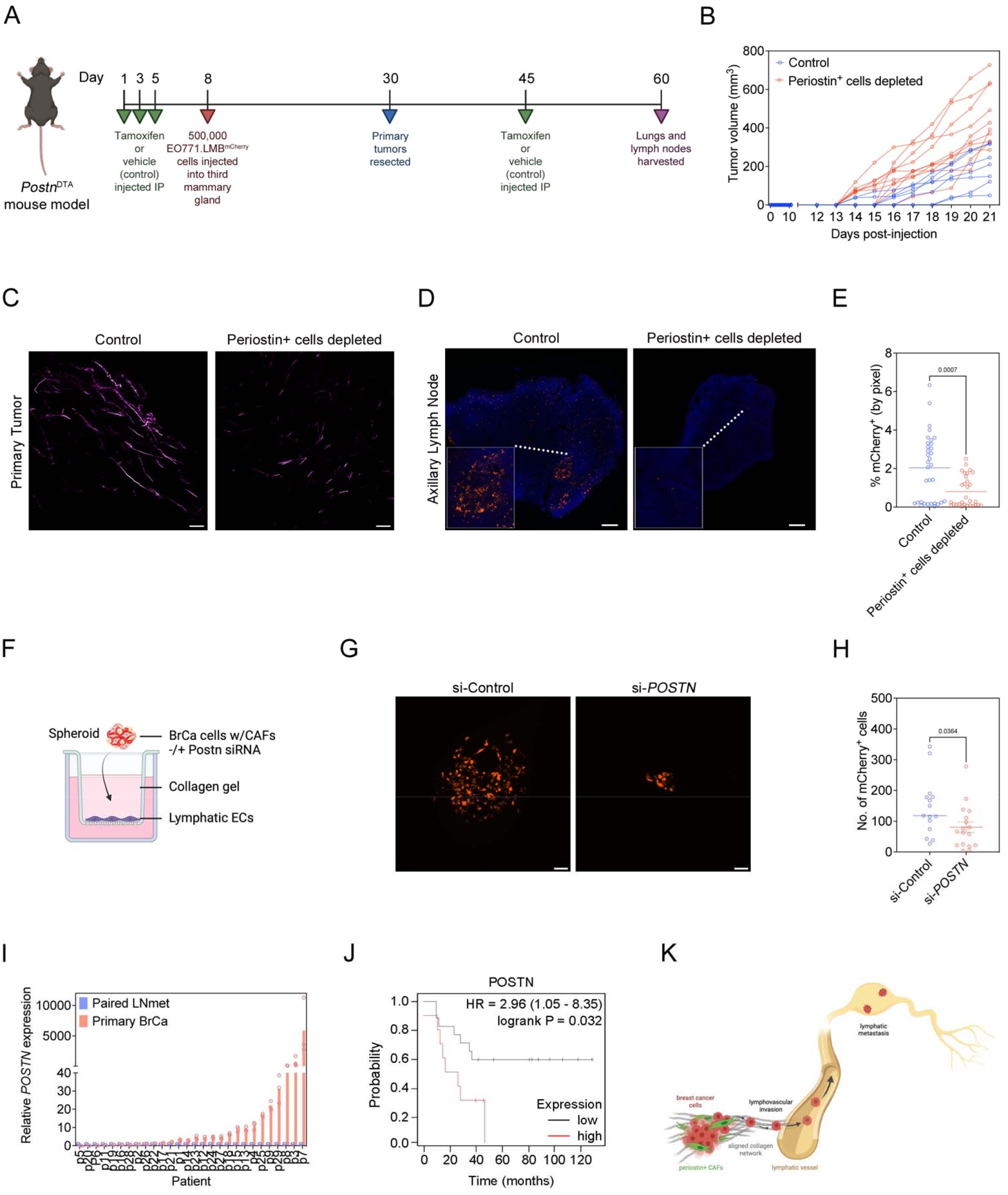
Periostin-expressing CAFs promote lymphatic metastasis by remodeling the extracellular matrix and directing lymphovascular invasion along organized collagen fibers. (A) Study design including tamoxifen (or vehicle-only control) treatment and orthotopic EO771.LMB mammary tumor injections in *Postn*^DTA^ mice. (B) Tumor volume measurements of highly metastatic EO771.LMB mammary tumors in control versus periostin+ cell-depleted mice. Each line represents an individual mouse (n = 8-10 mice per group). (C) Second harmonic generation (SHG) images of intratumoral collagen fibers (pseudo-colored in purple) in control versus periostin^+^ cell-depleted mice. Scale bars: 150 μm. (D) Tissue tilescans of axillary lymph nodes from control versus periostin^+^ cell-depleted mice bearing highly metastatic EO771.LMB mammary tumors. Tumor cells labelled with mCherry and nuclei counterstained with DAPI. Scale bars: 500 μm, insets are 3x zoom. (E) Percentage of tissue area positive for mCherry in serial sections of axillary lymph nodes from control versus periostin^+^ cell-depleted mice bearing highly metastatic EO771.LMB mammary tumors. Each data point in the scatter plot represents an individual histological section (n = 8-10 mice per group). Statistics shown for unpaired Student’s t test. (F) Experimental setup for in vitro lymphovascular invasion assay using MDA-MB-231^mCherry^ human breast cancer cells, primary human breast CAFs, and primary human lymphatic endothelial cells. (G) Representative fluorescent images of transmigrated MDA-MB-231^mCherry^ tumor cells that have invaded across the lymphatic endothelial cell barrier in the in vitro lymphovascular invasion assay. Breast cancer cells were co-cultured in spheroids with primary human breast CAFs that were pre-treated with either non-targeting control siRNA (si-Control) or periostin-targeting siRNA (si*-POSTN)*. Scale bar: 100 μm. (H) Quantification of the number of MDA-MB-231^mCherry^ tumor cells that have invaded across the lymphatic endothelial cell barrier in the in vitro lymphovascular invasion assay. Each data point represents an individual spheroid. Experiment performed in triplicate. Statistics shown for unpaired Student’s t test. (I) RT-qPCR analysis of periostin expression in paired primary breast cancer specimens (Primary BrCa) and lymph node metastases (Paired LNmet) from human breast cancer patients (n = 28 patients). qPCR performed in triplicate. (J) Kaplan-Meier plot of overall survival in lymph node positive breast cancer patients stratified based on periostin protein expression (n = 27 patients). (K) Summary model of how periostin-expressing CAFs promote lymphatic metastasis by remodeling the extracellular matrix and directing lymphovascular invasion along collagen fibers.

Surprisingly, depleting periostin-expressing cells accelerated the growth of primary mammary tumors which would indicate a growth-restraining feature of periostin-expressing cells at sites of primary growth by breast cancer cells (Figure 7B). In line with our previous observations, mammary tumors in which periostin-expressing cells had been depleted displayed impaired intratumoral collagen organization as detected by SHG imaging (Figure 7C), as collagen fibers were less aligned (Supplemental Figure 5A) and shorter in the tumors of periostin^+^ cell-depleted mice (235.0 ± 8.0 µm versus 154.9 ± 5.5 µm) (Supplemental Figure 5B). Periostin^+^ cell depletion also dramatically reduced lymphatic metastasis of highly metastatic EO771.LMB^mCherry^ mammary cancer cells. The metastatic burden in the draining axillary lymph nodes of periostin^+^ cell-depleted mice was reduced ∼3-fold (2% ± 0.3% versus 0.8% ± 0.1%) (Figure 7D,E), suggesting that periostin-expressing cells might enable lymphatic metastasis. Consistent with our previous results, very few EO771.LMB^mCherry^ mammary cancer cells reached the lungs in both the vehicle-control mice and periostin^+^ cell-depleted mice, and we did not observe a statistically significant reduction in pulmonary metastases following periostin^+^ cell depletion although the percentage of tissue area positive for mCherry^+^ cancer cells was trending downward in the periostin^+^ cell-depleted mice (Supplemental Figure 5C,D). Since targeting periostin on the cellular level reduced lymphatic metastasis, we then asked whether attenuating periostin expression on a molecular level would have a similar effect. Periostin expression in mouse mammary fibroblasts is regulated by TGFβ (Supplemental Figure 6A), so we hypothesized that reducing TGFβ-mediated periostin activation by conditionally knocking out a receptor in the TGFβ pathway (*Tgfbr2)* in periostin-expressing cells would similarly reduce lymphatic metastasis. Using *Postn-Cre:Tgfbr2*^*fl/fl*^ mice, we found a significant reduction in metastasis of highly-metastatic EO771. LMB^mCherry^ mammary cancer cells to the draining axillary lymph node following *Tgfbr2* knockout (Supplemental Figure 6A,B). Given the reduction in lymphatic metastasis following periostin depletion in vivo, we then modelled lymphovascular invasion in vitro to determine whether there is a role for periostin^+^ CAFs in guiding the intravasation of cancer cells across the lymphatic endothelial cell barrier. Using a modified spheroid/transwell migration assay (Figure 7F), we found that treating primary human breast CAFs with periostin-targeting siRNA inhibited their ability to promote the invasion of MDA-MB-231^mCherry^ human breast cancer cells through a collagen matrix and across a primary human lymphatic endothelial cell barrier (Figure 7G). Following periostin knockdown in CAFs, significantly fewer mCherry-labelled cancer cells crossed the lymphatic endothelial cell barrier compared to control (144 ± 24 cells versus 81 ± 17 cells) (Figure 7H), indicating that periostin-expressing CAFs promote lymphovascular invasion of breast cancer cells.

Taken together, our in vivo depletion studies and in vitro functional assays reveal a role for periostin-expressing CAFs in driving collagen-mediated lymphovascular invasion of cancer cells, resulting in lymphatic metastasis to the draining lymph node. To assess whether this pro-metastatic role for periostin within the primary tumor is reflected in clinical specimens, we used RT-qPCR analysis to measure relative periostin expression in paired primary breast tumors and lymph node metastases from 28 metastatic breast cancer patients obtained from the UNC Chapel Hill tissue biorepository. We found that periostin expression was higher in the primary tumors compared to their paired lymph node samples in 68% of patients (Figure 7I), which is consistent with periostin functioning within the primary TME to promote breast cancer cell escape and metastasis. To determine if periostin abundance is associated with poor patient outcome, we used the Kaplan-Meier Plotter to perform survival analysis of lymph node positive breast cancer patients stratified based on high or low periostin protein expression (Figure 7J). As predicted, periostin expression significantly correlated with decreased overall survival probability in this patient cohort, further supporting a clinically-relevant role for periostin in driving disease progression (Figure 7K).

## Discussion

Considerable progress has been made in the field of CAF biology using single-cell sequencing technologies to identify molecularly diverse CAF subpopulations. While expression profiles can indicate potential functional roles for these subclasses of CAFs, the clustering algorithms used to analyze sequencing data can yield subpopulations that are not necessarily spatially or pathologically relevant. Therefore, there remains a need to track these cells in situ and validate the functions of CAF subpopulations both in vitro and in vivo to determine their biological importance during tumor progression. In the present study, we have used lineage tracing strategies to functionally characterize a population of cells marked by expression of periostin – a matricellular protein associated with metastasis and expressed by the desmoplastic stroma of human breast cancers. We show that periostin is expressed by CAFs and perivascular-like cells and is enriched in advanced stage and lymph node positive human breast cancer samples. Our work also identifies a population of periostin-expressing PVL-CAFs that are enriched at the lymphatic vessel periphery and are differentially activated in highly-metastatic mammary tumors compared to their low-metastatic counterparts. In addition to quantitative differences, we observe phenotypic variation in the periostin-expressing cells between tumor types. The ZSGreen-labelled periostin-expressing cells in highly-metastatic tumors are larger and resemble the classical stellate shape of activated myofibroblasts whereas the periostin-expressing cells in low-metastatic tumors are smaller and rounded, reflecting the different activation states between the two populations. Together, this shows that the metastatic potential of cancer cells informs the activation of periostin-expressing cells in the primary tumor microenvironment. Interestingly, we find that the same is true at metastatic and premetastatic sites. We show that periostin-expressing cells are more abundant in the draining lymph nodes, lungs, and contralateral mammary glands of mice bearing highly-metastatic mammary tumors compared to those implanted with low-metastatic mammary tumors. A limitation of our spontaneous metastasis model is that relatively few mammary cancer cells reach the lungs before primary tumors re-grow to the maximum volume at which mice must be sacrificed. In order to address this, we used an experimental model of metastasis in which mammary cancer cells were injected into the tail vein of lineage tracing mice. Both low- and highly-metastatic mammary cancer lines established macrometastases in the lungs but marginally activated periostin-expressing cells despite a greater number of cancer cells present in these tissues. This indicates that periostin activation in a primary tumor is critical for robust activation of periostin at secondary sites rather than activation occurring at the secondary site once cancer cells have already colonized the tissue. These findings support previous reports that periostin is activated in premetastatic (24) and metastatic niches (26) where it may function to promote cancer cell survival and outgrowth, though our study focuses on an earlier pro-metastatic role for periostin within the primary tumor.

We find that periostin-expressing cell abundance is associated with intratumoral collagen organization, with straighter, longer, more continuous, and more aligned collagen fibers in highly-metastatic (periostin^high^) tumors. Our in vitro functional assays reveal that this difference in collagen matrix architecture is due to differences in intratumoral periostin abundance, as knocking down periostin expression in primary human breast CAFs reduces expression of a number of matrix-related proteins including collagens. This reduction in collagen secretion leads to a deficient and unorganized matrix, which impairs the ability of primary human breast CAFs to engage the matrix, spread, and migrate. These spreading and migratory defects are rescued by treatment with recombinant human periostin, suggesting that the differences in periostin-expressing cell morphology we observe in vivo in low-versus highly-metastatic tumors are directly related to differences in intratumoral periostin levels. We also show that periostin knockdown selectively inhibits collective cell invasion of primary human breast CAFs and breast cancer cells through collagen matrices. Though the timing of our in vitro assay is too short for CAFs to remodel collagen into the highly organized fibrillar networks we observe in periostin^high^ tumors in vivo, our results reflect that the inhibitory effect of periostin knockdown on early steps of collagen remodeling is sufficient to reduce collective cell invasion. Additionally, we can’t rule out that proliferation of the human breast cancer cells could contribute to the differences in the area of invasion that we observed, although we used low serum conditions and a 24-hour timepoint to minimize this effect. If CAF-derived periostin promotes collagen cross-linking as proposed, then it’s possible that cancer cells could engage the resulting matrix differently in control versus periostin-knockdown spheroids. This would potentially increase cancer cell proliferation in control spheroids as there is an established role for stiffened ECM in driving tumor proliferation (35), though we observe the opposite in vivo with depletion of periostin-expressing cells leading to a slightly increased primary tumor growth despite impaired collagen organization. This accelerated tumor growth following periostin^+^ cell depletion could be attributed to the loss of an organized collagen matrix, as there is evidence that fibrillar collagen can mechanically restrain tumor growth (37). Alternatively, it is possible that periostin-expressing cell depletion enhances primary tumor growth by shifting the immune landscape within tumors as has been previously observed upon broad CAF depletion in a model of pancreatic cancer (13). Therefore, an important future direction is to investigate the immune populations present in control versus periostin^+^ cell-depleted mammary tumors to determine if periostin-expressing CAFs perform immune-modulatory functions in addition to their roles in matrix remodeling.

Although our lineage tracing studies reveal that periostin-expressing cells make up a relatively small proportion of the total tumor tissue, they have a dramatic effect on the matrix architecture of the tumor microenvironment and lymphatic metastasis as evidenced by our periostin^+^ cell depletion tumor study. Our data reveal that periostin-expressing PVL-CAFs are instrumental in mediating lymphatic metastasis by depositing an organized collagen matrix and promoting lymphovascular invasion of breast cancer cells. CAFs and collagen alignment have been shown to play a role in hematogenous metastasis (38-41), but our work shows that periostin-expressing CAFs selectively function in promoting lymphatic metastasis, as their depletion markedly reduces metastatic burden in the lymph node but does not significantly affect pulmonary metastasis. These findings agree with and build upon recently published work showing that CAF-derived periostin can mediate metastasis by promoting lymphatic vessel permeability in an experimental model of popliteal lymph node metastasis (41), though this work used gain of function experiments to determine the effect of injected periostin on lymphatic metastasis of cervical squamous cell carcinoma whereas we have used genetic tools to determine the function of periostin^+^ cells in spontaneous breast cancer metastasis. Similarly, another study of periostin in experimental lymph node metastasis has shown an association between periostin deposition and lymphangiogenesis within lymph nodes prior to colonization by murine melanoma cells (42). This finding is consistent with our lineage tracing results showing periostin activation in lymph nodes as part of pre-metastatic niche formation, though our work emphasizes an ECM-remodeling role for periostin within the primary tumor site. Our mouse model recapitulates the desmoplastic stromal reaction linked to poor prognosis of breast cancer patients and allows us to deplete the periostin^+^ CAFs that contribute to this desmoplasia in situ to reveal their critical role in collagen-mediated lymphovascular invasion. As our study is the first to lineage trace and deplete periostin-expressing cells in the tumor context, this strategy can and should be adapted to other orthotopic tumor models to reveal whether this CAF population is present and shares a similar role in promoting the lymphatic metastasis of other cancers, especially those also characterized by desmoplasia such as pancreatic and lung cancer (19, 43, 44).

Densely aligned fibrillar collagen distinguishes progressive breast tumors from in situ lesions and is associated with increased periostin deposition in human patients (27), so early targeting or re-programming of the periostin^+^ CAF population could be an intervention to help prevent breast tumors from advancing to invasive disease. An important therapeutic consideration would be the timing of targeting periostin^+^ cells, as we observe increased primary tumor growth rate following periostin^+^ cell depletion similar to the paradoxical results of other CAF depletion studies (12, 13). To avoid this effect and given that periostin is activated by tissue injury and inflammation, blocking periostin^+^ CAF activation may be most effective around the time of surgical resection of the tumor, though further preclinical studies are required to examine the effect of periostin blockade at different timepoints during tumor progression and whether this could be an effective strategy to normalize the ECM and attenuate lymphatic metastasis. The ability of desmoplasia to drive breast cancer metastasis by promoting a mesenchymal phenotype in cancer cells to support cancer cell outgrowth at secondary sites has been established (38, 45), but attempts to target mediators of this process have been primarily preclinical with limited and largely unsuccessful clinical trials (46-48). Our study identifies a population of CAFs, most likely PVL-CAFs, responsible for the collagen remodeling that drives desmoplasia and demonstrates their role in promoting lymphovascular invasion and lymphatic metastasis, revealing a potential new avenue for therapeutic intervention in the metastatic cascade.

## Materials and Methods

### In vivo animal studies

All tumor studies were performed in 8- to 12-week old female mice with a C57BL/6 genetic background. *Postn*^iZSGreen^ lineage tracing mice were generated by crossing *Postn*^MCM^ mice (Jax Stock No. 029645) with *ZSGreen*^l/s/l^ mice (Jax Stock No. 007906). *Postn*^iZSGreen^ mice were injected intraperitoneally (IP) with 75 mg/kg tamoxifen 3 times over the course of 7 days to induce ZSGreen labelling of periostin-expressing cells. In the low-versus highly-metastatic tumor studies, one million EO771^mCherry^ or EO771.LMB^mCherry^ cancer cells were orthotopically injected into the third mammary fat pad of tamoxifen-induced heterozygous *Postn*^*i*ZSGreen^ lineage tracing mice. Tumor volumes were measured with calipers, and tumors were surgically removed when they averaged ∼400 mm^3^. Mice were euthanized and mammary glands, lymph nodes, and lungs were harvested ∼2 weeks following tumor resection surgery. For the tail vein model of experimental metastasis, tamoxifen-induced *Postn*^iZSGreen^ mice were injected with one million EO771^mCherry^ or EO771.LMB^mCherry^ cancer cells into their tail vein. Mice were sacrificed and lungs were harvested 2 weeks following tumor cell injection. *Postn*^iZSGreen^ lineage tracing mice were crossed with ROSA-DTA mice (Jax Stock No. 009669) to generate *Postn*^DTA^ mice that allow for selective depletion of periostin-expressing cells. *Postn*^DTA^ mice were injected IP with 75 mg/kg tamoxifen 3 times over the course of 7 days to induce depletion of periostin-expressing cells. 500,000 mCherry-labelled EO771.LMB cells were orthotopically injected into the third mammary fat pad of heterozygous tamoxifen-induced (or vehicle-injected control) mice. Tumor volumes were measured with calipers, and tumors were surgically removed when they averaged ∼400 mm^3^. Mice were euthanized and mammary glands, lymph nodes, and lungs were harvested 4 weeks following tumor resection surgery. *Postn-Cre:Tgfbr2*^*fl/fl*^ mice were generated by crossing *Postn*^iZSGreen^ mice with *Tgfbr2*^*fl/fl*^ mice (Jax Stock No. 012603). *Postn-Cre:Tgfbr2*^*fl/fl*^ mice were injected IP with 75 mg/kg tamoxifen 3 times over the course of 7 days to induce *Tgfbr2* knockout in periostin-expressing cells. 500,000 mCherry-labelled EO771.LMB cells were orthotopically injected into the third mammary fat pad of tamoxifen-induced *Postn-Cre:Tgfbr2*^*fl/fl*^ mice or *Postn*^iZSGreen^ control mice. Tumor volumes were measured with calipers, and tumors were surgically removed when they averaged ∼400 mm^3^. Mice were euthanized and lymph nodes were harvested 3 weeks following tumor resection surgery. All experiments were performed in accordance with the University of Virginia guidelines for animal handling and care.

### Immunohistology

Resected tissues were fixed overnight in 4% PFA/PBS and moved to a 30% sucrose solution for an additional 24 hours before embedding in OCT. Cryosections were DAPI stained and mounted in Vectashield (Vector Labs H-1700-10), then imaged using a Nikon Eclipse Ti-E inverted microscope and NIS-Elements software. Fluorescent populations were quantified using the thresholding function in FIJI (Supplemental Figure 1).

### Cell lines, cell culture, and media

EO771 cells were obtained from CH3 Biostystems (940001) and transfected with mCherry. EO771.LMB cells were a gift from Dr. Chad Pecot at UNC Chapel Hill and transfected with mCherry. EO771^mCherry^ and EO771.LMB^mCherry^ cells were cultured in 4.5 g/l D-glucose DMEM (HG-DMEM) with 10% FBS. PyVMT cells were a gift from Dr. Melanie Rutkowski at UVA and cultured in RPMI1640 with 2mM L-Glutamine, 10% FBS, 0.5% sodium pyruvate, and 0.09% β-mercaptoethanol. MDA-MB-231 human breast cancer cells were purchased from ATCC (HTB-26) and transfected with mCherry. MDA-MB-231^mCherry^ cells were cultured in HG-DMEM with 10% FBS. Primary human breast CAFs were isolated and gifted to us by Dr. Melissa Troester at UNC Chapel Hill (49) and were cultured on 0.5% gelatin-coated plates in HG-DMEM with 20% FBS and 10 ng/mL bFGF. Primary human lymphatic endothelial cells (Cell Biologics H-6092) were cultured in Complete Human Endothelial Cell Medium (Cell Biologics H1168) on 0.5% gelatin-coated plates. Primary mouse mammary fibroblasts were isolated from tumor-naïve mice and cultured on 0.5% gelatin-coated plates in HG-DMEM with 20% FBS and 10 ng/mL bFGF. All cell media included Plasmocin (Invivogen ant-mpt) to prevent mycoplasma growth and antibiotics/antimycotics to prevent bacterial and fungal contamination. Cells were maintained at 37° C in 5% CO_2_ plus 20% O_2_.

### Breast tumor tissue microarray analysis

Antigen-retrieval was performed on a tissue microarray of paraffin-embedded breast tumor cores (US Biomax BR20829) at 95° C for 20 minutes. The microarray slide was rinsed in DI water then incubated with blocking buffer (5% BSA + 5% goat serum in TBS) for 1 hour at room temperature. The slide was then incubated with rabbit anti-periostin antibody (Abcam ab215199, 1:1000) overnight at 4° C. The next day, the slide was rinsed 3 times with TBS and incubated for 30 minutes at room temperature with Biocare’s MACH 3 Rabbit Probe (M3R533 G). Following another three TBS washes, the slide was incubated for 30 minutes at room temperature with Biocare’s MACH 3 Rabbit AP-Polymer (M3R533 G). The slide was then rinsed in TBS and developed using Biocare’s Warp Red Chromogen Kit (WR 806 H) for 8 minutes at room temperature. The slide was washed in TBS, counterstained for 3 seconds with hematoxylin QS, then rinsed in water and TBS. Finally, the microarray slide was dipped in 100% ethanol then xylene and mounted using Biocare’s Ecomount (EM897L). Fluorescent AP signal was imaged using a Nikon Eclipse Ti-E inverted microscope and NIS-Elements software and percent periostin positive area (by pixel) was quantified by thresholding in FIJI.

### Immunoblotting

Collection of cell lysates and Western blotting were carried out as previously described using standard methods (50). For analysis of secreted proteins, conditioned media was collected and concentrated 10X using Microsep Advance Centrifugal Filters (Pall Laboratory MCP010C41). Samples probed for collagen I were prepared and run under non-reducing conditions in a 7.5% gel. Primary antibodies: rabbit anti-periostin (Abcam ab14041, 1:1000), rabbit anti-collagen I (Abcam ab34710, 1:1000). Secondary HRP-conjugated antibody: peroxidase goat anti-rabbit IgG (Vector Laboratories PI-1000, 1:10,000).

### Real-time quantitative PCR (RT-qPCR)

Total RNA isolation was performed using a Quick-RNA Miniprep Kit (Zymo Research R1055) according to the manufacturer’s instructions. cDNA synthesis was completed using an iScript cDNA Synthesis Kit (Bio-Rad 1708891EDU), and qPCR reactions were carried out using a QuantStudio 12K Flex Real-time PCR System.

### Immunofluorescence

Fixed frozen tissue sections were stained using the following antibodies and concentrations: Cy3-SMA (TCF C6198, 1:100), CD31 (BD Biosciences 550274, 1:50), F4/80 (BioRad MCA497, 1:50), CK14 (Biolegend 905301, 1:250), Pdpn (R&D Systems AF3244, 1:200). Antibodies were diluted in blocking buffer with 5% BSA, 5% goat serum, and 1% Triton X-100. Primary incubations were performed overnight at 4° C and secondary incubations were performed for 1 hour at room temperature. Nuclei were counterstained with DAPI and slides were mounted using Vectashield (Vector Labs H-1700-10). Slides were imaged using a Nikon Eclipse Ti-E inverted microscope and NIS-Elements software.

### GeoMx Digital Spatial Profiling (DSP)

NanoString GeoMx DSP was used to quantify transcript numbers in spatially distinct populations of periostin-expressing cells in the naïve mammary glands of *Postn*^iZSGreen^ lineage tracing mice. Fixed frozen mammary glands were incubated with the NanoString mouse Whole Transcriptome Atlas (WTA) panel probes overnight, then stained with Cy3-SMA (TCF C6198), Pdpn (R&D Systems AF3244;Cy5 secondary), and the DNA stain Syto83 (Thermo S11364) as morphology markers to visualize tissue architecture. Stained slides were loaded onto the GeoMx instrument and scanned for region of interest (ROI) selection. Due to the filters in the optical system of the DSP platform, ZSGreen^+^ cells appear blue, SMA staining appears in green, and Pdpn staining appears in red in the ROIs. ROIs of ZSGreen^+^ cells near Pdpn^+^ structures were selected to represent lymphatic-adjacent populations, and ROIs of ZSGreen^+^ cells near SMA^+^ structures were selected to represent duct- and blood vessel-adjacent populations. After UV illumination of the ROIs, the eluent was collected via microcapillary aspiration and transferred into individual wells of a microtiter plate. The collected aspirates in the microtiter plate were then transferred to a PCR plate for library prep with Seq Code primers. PCR products were pooled and purified, then subjected to Illumina NGS Sequencing (Next Seq 2000). FASTQ sequencing files obtained from the NGS run were processed into digital count conversion (DCC) files by NanoString’s GeoMx NGS Pipeline software. The DCC files were then uploaded onto the GeoMx DSP, and quality control checks and differential gene expression analysis were performed.

### Second harmonic generation

Fixed frozen tumor sections were imaged using a Zeiss 710 Multiphoton Confocal microscope and collagen fiber length measurements were performed in FIJI. The FIJI plug-ins TWOMBLI (51) and OrientationJ (52) were used to analyze the number of endpoints, curvature (measured as mean change in angle along fibers), and directionality of the collagen fibers.

### siRNA transfection

For in vitro periostin knockdown studies, primary human breast CAFs were transfected with 75 nm human periostin-targeting stealth siRNA (si*-POSTN)* (ThermoFisher siRNA ID: HSS116400) or 75 nm non-targeting control siRNA (si-Control) (DHARMACON D-001810-02-05) for 5 hours, then media was replaced. This siRNA transfection was repeated after 24 hours, and media was replaced with low-serum media. Cells were then used for functional assays.

### Bulk RNAseq and analysis

Primary human breast CAFs were transfected with siRNA in quadruplicate as described above. RNA samples were harvested from cells and sent to Novogene for bulk RNA sequencing. Sequencing analysis was performed by the UVA Bioinformatics Core. On average we received 30 million paired ends for each of the replicates. RNAseq libraries were checked for their quality using the fastqc program (http://www.bioinformatics.babraham.ac.uk/projects/fastqc/). The results from fastqc were aggregated using multiqc software. In house developed programs was used for adaptor identification, and any contamination of adaptor sequence was removed with cutadapt (https://cutadapt.readthedocs.io/en/stable/). Reads were mapped with the “splice aware” aligner ‘STAR’ to the transcriptome and genome of mm10 genome build. The HTseq software was used to count aligned reads that map onto each gene. The count table was imported into R to perform differential gene expression analysis using the DESeq2 package. Low expressed genes (genes expressed only in a few replicates and had low counts) were excluded from the analysis before identifying differentially expressed genes. Data normalization, dispersion estimates, and model fitting (negative binomial) were carried out with the DESeq function. The log-transformed, normalized gene expression of 500 most variable genes was used to perform an unsupervised principal component analysis. The differentially expressed genes were ranked based on the log2fold change and FDR corrected p-values. The ranked file was used to perform pathway analysis using GSEA software. The enriched pathways were selected based on enrichment scores as well as normalized enrichment scores.

### ECM deposition assay

Primary human breast CAFs were seeded (30,000 cells per well) on a gelatin-coated chamber slide and transfected with siRNA as described above. 48 hours after the second siRNA treatment, cells were lysed using ammonium hydroxide-based extraction buffer. The deposited ECM was washed in PBS and fixed with 2% PFA for 20 minutes. The ECM was then rinsed in PBS and incubated in blocking buffer with 1% Triton for 30 minutes to reduce non-specific antibody binding. The ECM was then incubated with primary antibody (rabbit anti-collagen I ab34710, 1:200) for 1 hour at room temperature and secondary antibody (goat anti-rabbit IgG Alexa Fluor 594 1:100) for 1 hour at room temperature. Slides were mounted using Vectashield (Vector Labs H-1700-10) and imaged using a Nikon Eclipse Ti-E inverted microscope and NIS-Elements software. Collagen fiber length measurements were performed using FIJI.

### Cell spreading assay

Primary human breast CAFs were seeded (1 × 10^5^ cells/mL) in a 6-well gelatin coated dish and transfected with siRNA as described above. 24 hours following knockdown, CAFs were detached and re-seeded at 10,000 cells per well on a gelatin-coated chamber slide. The “rescue” wells were treated with 0.8 ug/mL recombinant human periostin (BioLegend 770506). After 24 hours, cells were fixed in 4% PFA and stained with Alexa Fluor 594 Phalloidin (ThermoFisher Scientific A12381). Cells were imaged using a Nikon Eclipse Ti-E inverted microscope and NIS-Elements software, and cell area measurements were performed in FIJI.

### Wound closure scratch assay

Primary human breast CAFs were seeded (1 × 10^5^ cells/mL) in a 6-well gelatin coated dish and transfected with siRNA as described above. The next day, a P1000 pipette tip was used to create a scratch down the middle of the cell monolayer in each well. Each well was washed once with PBS to removed detached cells and low serum media was then added. “Rescue” wells were treated with 0.8 ug/mL recombinant human periostin (BioLegend 770506). Scratches were imaged at 0 hours, 12 hours, and 24 hours using a Nikon Eclipse Ti-E inverted microscope and NIS-Elements software, and percent wound closure was calculated using area measurements in FIJI.

### 3D CAF/cancer cell spheroid invasion assay

Primary human breast CAFs were seeded (1 × 10^5^ cells/mL) in a 6-well gelatin coated dish and transfected with siRNA as described above. The next day, CAFs were co-cultured 1:1 with MDA-MB-231^mCherry^ human breast cancer cells in 20 uL hanging droplets containing 5% methylcellulose on the inverted lid of a 100 mm tissue culture dish containing PBS. The CAF/cancer cell co-cultures were incubated for 24 hours to allow aggregation of cells into a spheroid, as confirmed under light microscopy. The CAF/cancer cell spheroids were then embedded in either Matrigel (Corning 356237) or rat-tail collagen type I (Ibidi 50201) according to manufacturer’s instructions in a 96-well plate. Spheroids were imaged at 12 and 24 hours using a Nikon Eclipse Ti-E inverted microscope and NIS-Elements software, and invasive area was quantified using FIJI. Representative images of spheroids were taken using a Zeiss LSM 880 confocal microscope.

### Trans-well migration assay

Primary human breast CAFs were seeded (1.5 × 10^5^ cells/mL) in a 6-well gelatin coated dish and transfected with siRNA as described above. The next day, CAFs were co-cultured with MDA-MB-231^mCherry^ human breast cancer cells into spheroids as described above. Transwell membrane inserts (Fisher Scientific 07-200-150) were coated with 0.5% gelatin and seeded with 60,000 primary human lymphatic endothelial cells. After 24 hours, CAF/cancer cell spheroids were embedded in rat-tail collagen type I (Ibidi 50201) in the transwell membrane inserts above the monolayer of primary human lymphatic endothelial cells. Inserts were cleared with a cotton swab and washed in PBS after 24 hours, so that only cells that had migrated across the lymphatic endothelial cell barrier and reached the bottom surface of the transwell insert remained. These cells were fixed in 4% PFA for 10 minutes at room temperature and washed in PBS. Transwell inserts were placed on slides and imaged using a Nikon Eclipse Ti-E inverted microscope and NIS-Elements software. Trans-migrated cancer cells were counted in FIJI.

### Kaplan-Meier Analysis

Kaplan-Meier Plotter was used to generate survival curves for lymph node positive breast cancer patients from the Tang_2018 patient cohort that were stratified based on low versus high periostin protein expression.

### Statistical analysis

All data points are shown, and horizontal lines on graphs represent median values. Descriptive numerical values in the text are expressed as mean value ± standard error of the mean (SEM). All statistical analyses were performed using GraphPad Prism software and P values less than 0.05 were considered significant.

## Acknowledgments

JLN is currently supported by an NCI Ruth L. Kirschstein NRSA for Individual Predoctoral Fellows Award (F31CA247407-02) and previously received support from the NIH/NCI training grant T32CA009109. ACD is supported by grants from the American Cancer Society (129755-RSG-16-176-DDC), the National Institutes of Health/National Cancer Institute (2RO1 CA177875 and RO1 CA2558451), the Melanoma Research Alliance (ID612638), and funds from the Emily Couric Cancer Center at the University of Virginia. Portions of this research were supported by the NCI Cancer Center Support Grant 5P30CA044579 and by the UVA Genome Analysis and Technology Core (RRID:SCR_018883). Additional support was provided by The University of Virginia Flow Cytometry Core (RRID: SCR_017829). The Sony MA900 Cell Sorter was funded through the NIH S10 instrument program (S10 Grant Number 1S10OD028518-1). This work used the Zeiss 710 multiphoton confocal microscope and Zeiss LSM 880 confocal microscope in the UVA Advanced Microscopy Facility which is supported by the University of Virginia School of Medicine. We would like to thank Natalia Dworak, Dr. Stacey Criswell, and Dr. Adrian Halme for their assistance using this equipment. Illumina NGS sequencing for the spatial RNA profiling was performed by the University of Minnesota Genomics Center. We would also like to thank Marya Dunlap-Brown of the Molecular Immunologic and Translational Sciences (MITS) Core at UVA for her help with mouse surgeries. All mouse tissues were sectioned using the Thermo CryoStar NX50 cryostat in the UVA Biorepository and Tissue Research Facility (BRTF), and we wish to thank Angela Miller for her training and technical assistance. BioRender was used to design all schematics in the manuscript.

## Author contributions

JLN and ACD conceptualized the study and wrote the manuscript. JLN carried out the experiments. DJK and JVM assisted with in vitro functional assays. PP and JWF performed spatial RNA profiling, and PK performed bulk RNAseq data analysis. LE and CP provided qPCR analysis of *POSTN* expression in paired primary tumors and lymph node metastases from human breast cancer patients. All authors have been provided with a copy of the complete manuscript prior to submission.

## Data and materials availability

All bioinformatics data will be made available immediately upon publication

## References

1. A. J. Redig, S. S. McAllister, Breast cancer as a systemic disease: a view of metastasis. J Intern Med 274, 113–126 (2013).

2. R. A. A. Mohammed et al., Improved Methods of Detection of Lymphovascular Invasion Demonstrate That It is the Predominant Method of Vascular Invasion in Breast Cancer and has Important Clinical Consequences. The American Journal of Surgical Pathology 31 (2007).

3. G. Houvenaeghel et al., Lymphovascular invasion has a significant prognostic impact in patients with early breast cancer, results from a large, national, multicenter, retrospective cohort study. ESMO Open 6 (2021).

4. E. A. Rakha et al., The prognostic significance of lymphovascular invasion in invasive breast carcinoma. Cancer 118, 3670–3680 (2012).

5. S. D. Nathanson, D. Kwon, A. Kapke, S. H. Alford, D. Chitale, The Role of Lymph Node Metastasis in the Systemic Dissemination of Breast Cancer. Annals of Surgical Oncology 16, 3396–3405 (2009).

6. R. Sabatier et al., Peritumoural vascular invasion: A major determinant of triple-negative breast cancer outcome. European Journal of Cancer 47, 1537–1545 (2011).

7. T. Liu, L. Zhou, D. Li, T. Andl, Y. Zhang, Cancer-Associated Fibroblasts Build and Secure the Tumor Microenvironment. Frontiers in Cell and Developmental Biology 7 (2019).

8. A. Labernadie et al., A mechanically active heterotypic E-cadherin/N-cadherin adhesion enables fibroblasts to drive cancer cell invasion. Nature Cell Biology 19, 224–237 (2017).

9. A. Marusyk et al., Spatial Proximity to Fibroblasts Impacts Molecular Features and Therapeutic Sensitivity of Breast Cancer Cells Influencing Clinical Outcomes. Cancer Res 76, 6495–6506 (2016).

10. B. Erdogan et al., Cancer-associated fibroblasts promote directional cancer cell migration by aligning fibronectin. J Cell Biol 216, 3799–3816 (2017).

11. E. Hirata et al., Intravital imaging reveals how BRAF inhibition generates drug-tolerant microenvironments with high integrin β1/FAK signaling. Cancer Cell 27, 574–588 (2015).

12. A. D. Rhim et al., Stromal elements act to restrain, rather than support, pancreatic ductal adenocarcinoma. Cancer Cell 25, 735–747 (2014).

13. B. C. Özdemir et al., Depletion of carcinoma-associated fibroblasts and fibrosis induces immunosuppression and accelerates pancreas cancer with reduced survival. Cancer Cell 25, 719–734 (2014).

14. E. J. Helms et al., Mesenchymal Lineage Heterogeneity Underlies Non-Redundant Functions of Pancreatic Cancer-Associated Fibroblasts. Cancer Discovery, candisc.0601.2021 (2021).

15. M. Bartoschek et al., Spatially and functionally distinct subclasses of breast cancer-associated fibroblasts revealed by single cell RNA sequencing. Nature Communications 9, 5150 (2018).

16. S. Davidson et al., Single-Cell RNA Sequencing Reveals a Dynamic Stromal Niche That Supports Tumor Growth. Cell Reports 31, 107628 (2020).

17. S. V. Puram et al., Single-Cell Transcriptomic Analysis of Primary and Metastatic Tumor Ecosystems in Head and Neck Cancer. Cell 171, 1611–1624.e1624 (2017).

18. G. Friedman et al., Cancer-associated fibroblast compositions change with breast cancer progression linking the ratio of S100A4+ and PDPN+ CAFs to clinical outcome. Nature Cancer 1, 692–708 (2020).

19. D. Lambrechts et al., Phenotype molding of stromal cells in the lung tumor microenvironment. Nat Med 24, 1277–1289 (2018).

20. S. Chen et al., Single-cell analysis reveals transcriptomic remodellings in distinct cell types that contribute to human prostate cancer progression. Nature Cell Biology 23, 87–98 (2021).

21. D. Öhlund et al., Distinct populations of inflammatory fibroblasts and myofibroblasts in pancreatic cancer. J Exp Med 214, 579–596 (2017).

22. S. Z. Wu et al., Stromal cell diversity associated with immune evasion in human triple-negative breast cancer. The EMBO Journal 39, e104063 (2020).

23. T. Maruhashi, I. Kii, M. Saito, A. Kudo, Interaction between periostin and BMP-1 promotes proteolytic activation of lysyl oxidase. J Biol Chem 285, 13294–13303 (2010).

24. Z. Wang et al., Periostin promotes immunosuppressive premetastatic niche formation to facilitate breast tumour metastasis. J Pathol 239, 484–495 (2016).

25. J. Soikkeli et al., Metastatic outgrowth encompasses COL-I, FN1, and POSTN up-regulation and assembly to fibrillar networks regulating cell adhesion, migration, and growth. Am J Pathol 177, 387–403 (2010).

26. I. Malanchi et al., Interactions between cancer stem cells and their niche govern metastatic colonization. Nature 481, 85–89 (2012).

27. T. Risom et al., Transition to invasive breast cancer is associated with progressive changes in the structure and composition of tumor stroma. Cell 185, 299–310.e218 (2022).

28. O. Kanisicak et al., Genetic lineage tracing defines myofibroblast origin and function in the injured heart. Nature Communications 7, 12260 (2016).

29. S. Z. Wu et al., A single-cell and spatially resolved atlas of human breast cancers. Nature Genetics 53, 1334–1347 (2021).

30. C. N. Johnstone et al., Functional and molecular characterisation of EO771.LMB tumours, a new C57BL/6-mouse-derived model of spontaneously metastatic mammary cancer. Dis Model Mech 8, 237–251 (2015).

31. S. Alkner et al., Contralateral breast cancer can represent a metastatic spread of the first primary tumor: determination of clonal relationship between contralateral breast cancers using next-generation whole genome sequencing. Breast Cancer Research 17, 102 (2015).

32. V. Vichapat et al., Patterns of metastasis in women with metachronous contralateral breast cancer. British Journal of Cancer 107, 221–223 (2012).

33. R. A. Norris et al., Periostin regulates collagen fibrillogenesis and the biomechanical properties of connective tissues. J Cell Biochem 101, 695–711 (2007).

34. P. P. Provenzano et al., Collagen reorganization at the tumor-stromal interface facilitates local invasion. BMC Medicine 4, 38 (2006).

35. K. R. Levental et al., Matrix crosslinking forces tumor progression by enhancing integrin signaling. Cell 139, 891–906 (2009).

36. M. W. Conklin et al., Aligned collagen is a prognostic signature for survival in human breast carcinoma. Am J Pathol 178, 1221–1232 (2011).

37. S. Bhattacharjee et al., Tumor restriction by type I collagen opposes tumor-promoting effects of cancer-associated fibroblasts. J Clin Invest 131 (2021).

38. N. Dumont et al., Breast fibroblasts modulate early dissemination, tumorigenesis, and metastasis through alteration of extracellular matrix characteristics. Neoplasia 15, 249–262 (2013).

39. J. Condeelis, J. E. Segall, Intravital imaging of cell movement in tumours. Nature Reviews Cancer 3, 921–930 (2003).

40. W. Wang et al., Single cell behavior in metastatic primary mammary tumors correlated with gene expression patterns revealed by molecular profiling. Cancer Res 62, 6278–6288 (2002).

41. W. Han et al., Oriented collagen fibers direct tumor cell intravasation. Proceedings of the National Academy of Sciences 113, 11208–11213 (2016).

42. L. Gillot et al., Periostin in lymph node pre-metastatic niches governs lymphatic endothelial cell functions and metastatic colonization. Cell Mol Life Sci 79, 295 (2022).

43. C. J. Whatcott et al., Desmoplasia in Primary Tumors and Metastatic Lesions of Pancreatic Cancer. Clinical Cancer Research 21, 3561–3568 (2015).

44. N. K. Altorki et al., The lung microenvironment: an important regulator of tumour growth and metastasis. Nature Reviews Cancer 19, 9–31 (2019).

45. T. R. Cox et al., LOX-mediated collagen crosslinking is responsible for fibrosis-enhanced metastasis. Cancer Res 73, 1721–1732 (2013).

46. S. Ferreira, N. Saraiva, P. Rijo, A. S. Fernandes, LOXL2 Inhibitors and Breast Cancer Progression. Antioxidants (Basel) 10 (2021).

47. T. Nandi, S. Pradyuth, A. K. Singh, D. Chitkara, A. Mittal, Therapeutic agents for targeting desmoplasia: current status and emerging trends. Drug Discovery Today 25, 2046–2055 (2020).

48. J. Huang et al., Extracellular matrix and its therapeutic potential for cancer treatment. Signal Transduction and Targeted Therapy 6, 153 (2021).

49. H. A. Brauer et al., Impact of Tumor Microenvironment and Epithelial Phenotypes on Metabolism in Breast Cancer. Clinical Cancer Research 19, 571–585 (2013).

50. L. Xiao et al., Tumor Endothelial Cells with Distinct Patterns of TGFβ-Driven Endothelial-to-Mesenchymal Transition. Cancer Research 75, 1244 (2015).

51. E. Wershof et al., A FIJI macro for quantifying pattern in extracellular matrix. Life Sci Alliance 4 (2021).

52. Z. Püspöki, M. Storath, D. Sage, M. Unser, Transforms and Operators for Directional Bioimage Analysis: A Survey. Adv Anat Embryol Cell Biol 219, 69–93 (2016).

