## Supplemental Figure 1 for "Periostin^+^ stromal cells guide lymphovascular invasion by cancer cells"

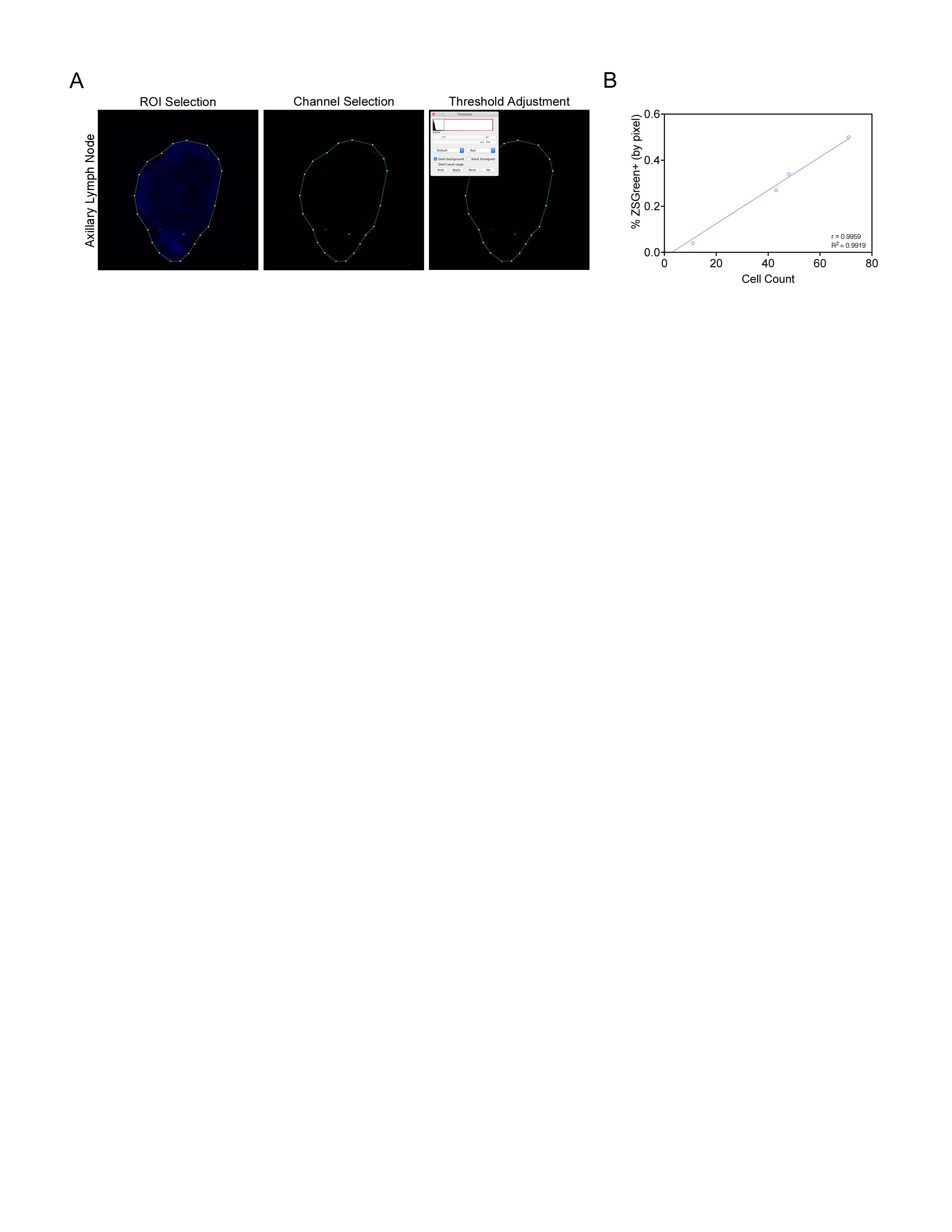

**Supplemental Figure 1. *Thresholding strategy for fluorescent area quantification.*** (A) Workflow in FIJI to quantify fluorescent area by thresholding. First, region of interest (ROI) is selected around the tissue boundary using DAPI nuclear counterstain as a guide so that background pixels are excluded from quantification. Next, fluorescent channel is selected to quantify population of interest. Finally, threshold is adjusted so that all fluorescent cells are captured with minimal noise. Positive areas are highlighted in red by thresholding tool, and percent positive measurement is displayed in Threshold window. (B) Correlation between individual cell count and percentage of tissue area positive for ZSGreen (by pixel) measured by thresholding. Pearson r and R squared values shown.

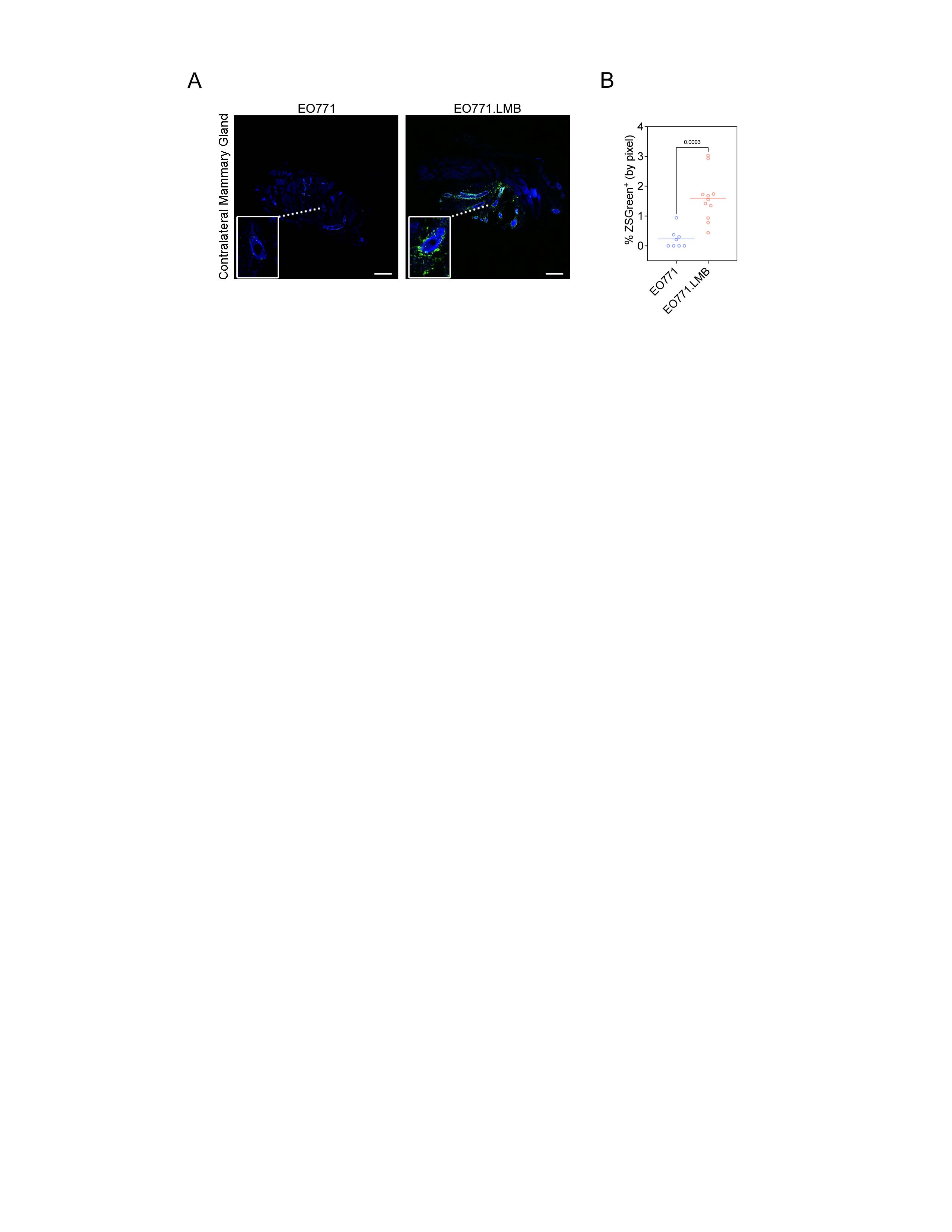

**Supplemental Figure 2. *Periostin-expressing PVL-CAFs are more abundant in the premetastatic niches of mice bearing highly metastatic mammary tumors.*** (A) Tissue tilescans of contralateral mammary glands from *Postn*^iZSGreen^ lineage tracing mice bearing low metastatic (EO771) or highly metastatic (EO771.LMB) mammary tumors. Periostin-expressing cells genetically labelled with ZSGreen, and tumor cells labelled with mCherry (not detected in these tissues). Nuclei counterstained with DAPI. Scale bars: 500 μm, insets are 3x zoom. (B) Percentage of tissue area positive for ZSGreen in serial sections of contralateral mammary glands of mice bearing EO771 or EO771.LMB mammary tumors. Each data point represents a different histological section (n = 4-5 mice per group). Statistics shown for unpaired Student’s t test.

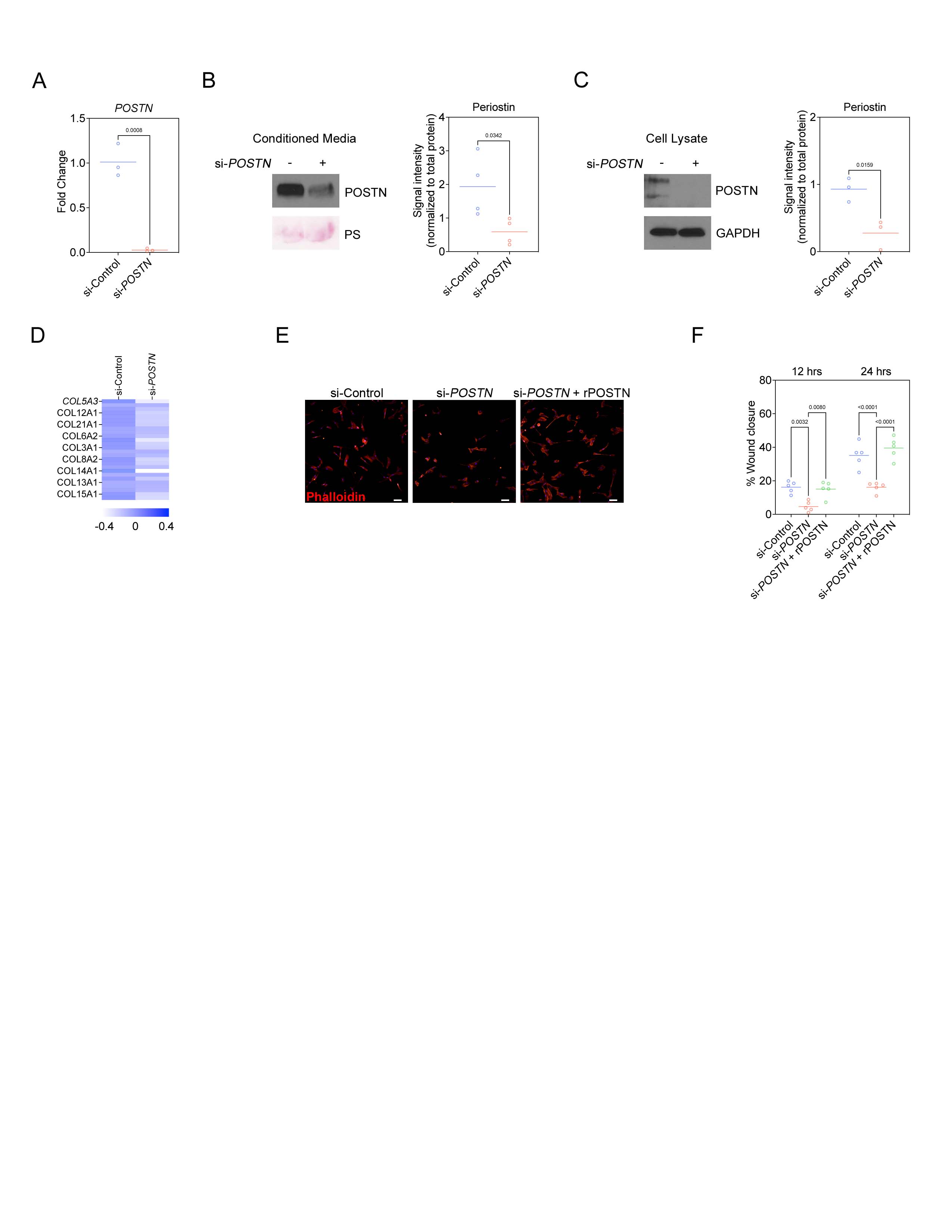

**Supplemental Figure 3. *Periostin knockdown in primary human breast CAFs inhibits cell spreading and migration.*** (A) qPCR analysis to confirm periostin expression is reduced in primary human breast CAFs treated with periostin-targeting siRNA (si-*POSTN*) compared to cells treated with a non-targeting control siRNA (si-Control). Performed in triplicate. Statistics shown for unpaired Student’s t test. (B) Western blot for secreted periostin in primary human breast CAFs treated with si-Control or si*-POSTN.* Ponceau stain (PS) shown as loading control. Protein signal intensity quantified on the right, performed in quadruplicate. Statistics shown for unpaired Student’s t test. (C) Western blot for intracellular periostin in primary human breast CAFs treated with si-Control or si*-POSTN.* GAPDH shown as loading control. Protein signal intensity quantified on the right, performed in triplicate. Statistics shown for unpaired Student’s t test. (D) Heat map of gene expression of collagen family proteins in primary human breast CAFs treated with si-Control or si*-POSTN.* (E) Immunofluorescence images of phalloidin staining of primary human breast CAFs treated with si-Control or si*-POSTN* ± recombinant human periostin (rPOSTN). Nuclei counterstained with DAPI. Scale bars: 100 μm. (F) Scratch assay quantification of percent wound closure over time by primary human breast CAFs treated with si-Control or si*-POSTN* ± recombinant human periostin (rPOSTN) to measure cell migration. Each data point represents a different scratch replicate. Statistics shown for 2-way ANOVA, Tukey’s multiple comparisons test.

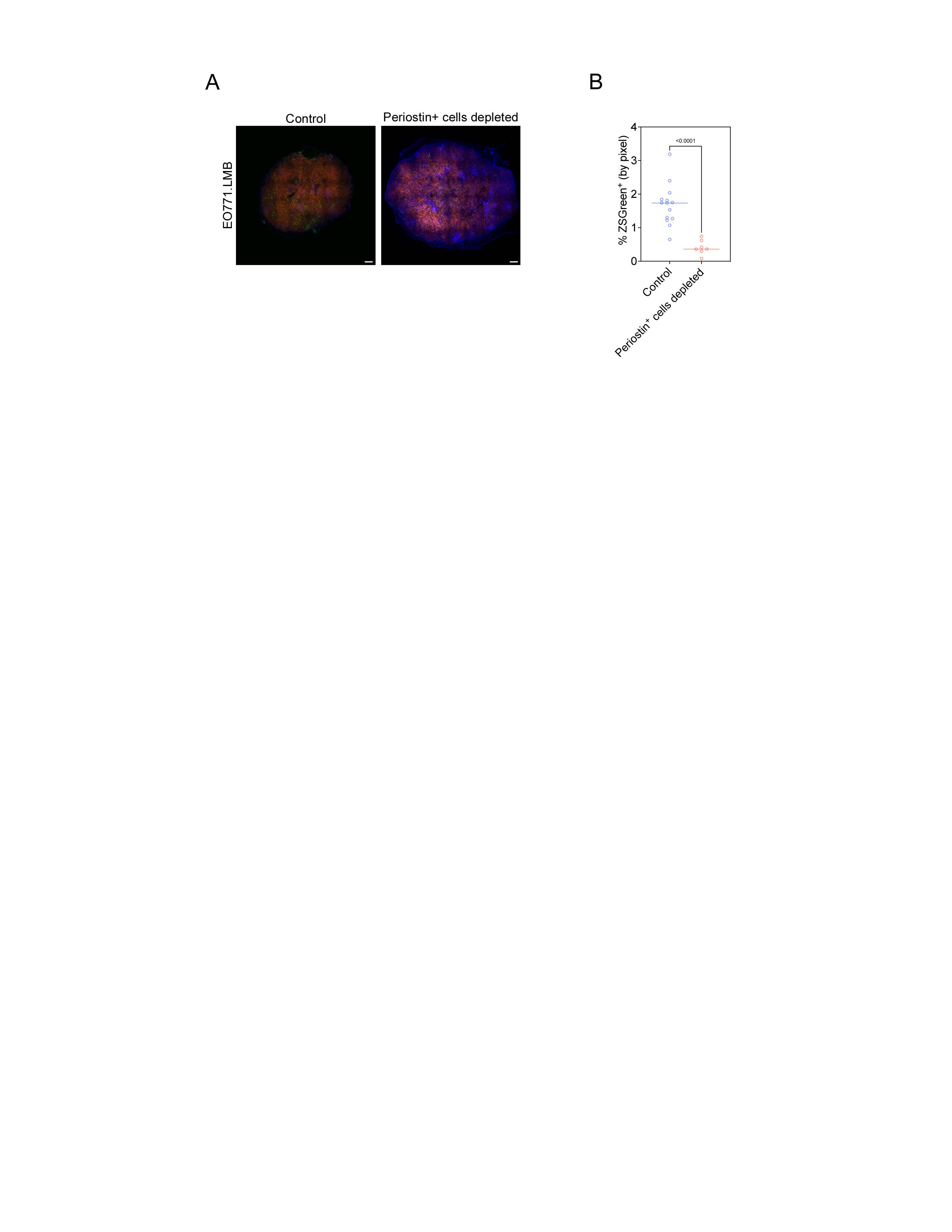

**Supplemental Figure 4. *Confirmation of periostin-expressing cell depletion in mammary tumors of Postn^DTA^ mice.*** (A) Tissue tilescans of EO771.LMB mammary tumors from *Postn*^iZSGreen^ control mice (left) and *Postn*^DTA^ mice (right). Tumor cells labelled with mCherry and periostin-expressing cells genetically labelled with ZSGreen. Nuclei counterstained with DAPI. Scale bars: 500 μm. (B) Percentage of tissue area positive for ZSGreen in EO771.LMB tumors (n = 3-5 mice per group). Each data point represents a different histological section. Statistics shown for unpaired Student’s t test.

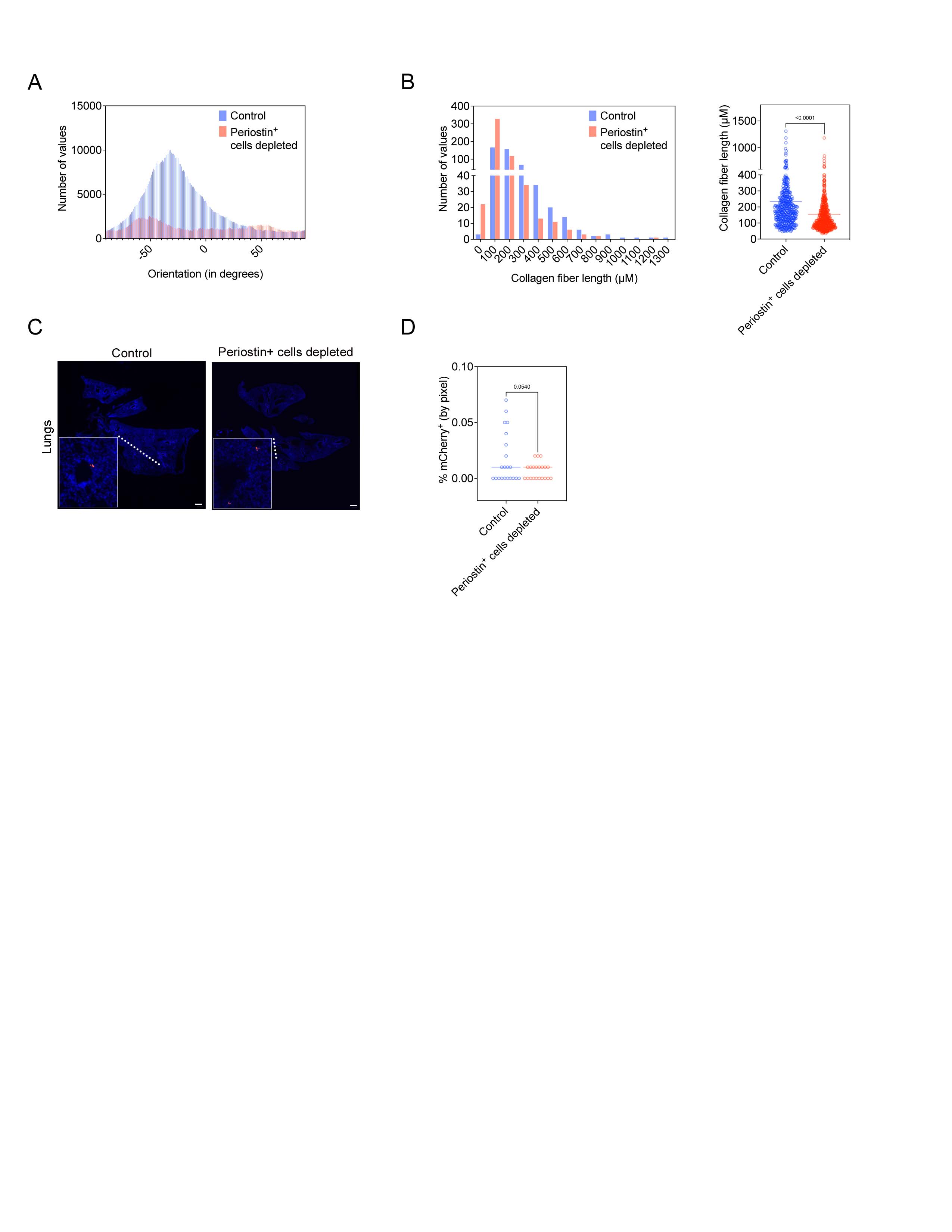

**Supplemental Figure 5. *Depleting periostin-expressing PVL-CAFs alters matrix architecture in primary tumors but does not significantly reduce metastatic burden in the lung.*** (A) Representative histogram of collagen fiber orientation in EO771.LMB tumors from vehicle-only injected control mice and periostin^+^ cell-depleted mice. A peaked histogram represents aligned fibers whereas a flat histogram represents random organization. (B) Quantification of collagen fiber length in EO771.LMB tumors from vehicle-only injected control mice and periostin^+^ cell-depleted mice, represented as a histogram (left) and scatter plot (right). Each point represents an individual fiber. Between 475 and 540 fibers quantified from multiple histological sections (n = 3-4 mice per group) and results compared using unpaired Student’s t test. (C) Tissue tilescans of lungs from either vehicle-only injected control mice or periostin^+^ cell-depleted mice bearing EO771.LMB mammary tumors. Tumor cells labelled with mCherry and nuclei counterstained with DAPI. Scale bars: 500 μm, insets are 6x zoom. (D) Percentage of tissue area positive for mCherry in serial sections of lungs from either vehicle-only injected control mice or periostin^+^ cell-depleted mice bearing EO771.LMB mammary tumors. Each point represents a different histological section (n = 7 mice per group). Statistics shown for unpaired Student’s t test.

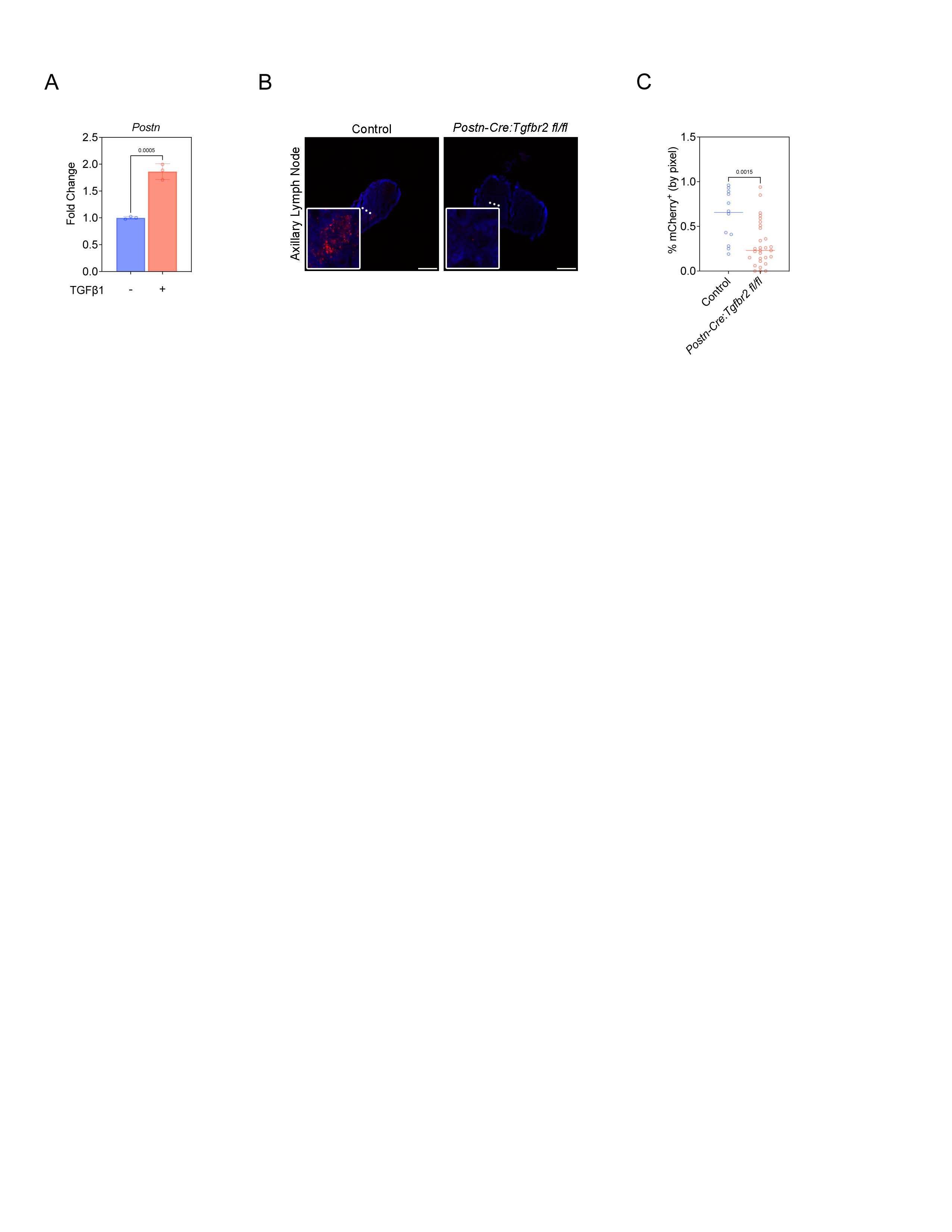

**Supplemental Figure 6. *Tgbfr2 knockout in periostin-expressing cells reduces lymphatic metastasis.*** (A) qPCR analysis of periostin expression in primary mouse mammary fibroblasts treated with 10 ng/mL TGFβ1. Statistics shown for unpaired Student’s t test. (B) Tissue tilescans of axillary lymph nodes from control mice or *Postn-Cre:Tgfbr2^fl/fl^* mice. Tumor cells labelled with mCherry and nuclei counterstained with DAPI. Scale bars: 500 μm, insets are 6x zoom. (C) Percentage of tissue area positive for mCherry in serial sections of axillary lymph nodes from either control mice or *Postn-Cre:Tgfbr2^fl/fl^* mice bearing EO771.LMB mammary tumors. Each point represents a different histological section (n = 4-10 mice per group). Statistics shown for unpaired Student’s t test.
